# Binding patterns of RNA binding proteins to repeat-derived RNA sequences reveal putative functional RNA elements

**DOI:** 10.1101/2021.06.03.446901

**Authors:** Masahiro Onoguchi, Chao Zeng, Ayako Matsumaru, Michiaki Hamada

## Abstract

Recent reports have revealed that repeat-derived sequences embedded in introns or long noncoding RNAs (lncRNAs) are targets of RNA binding proteins (RBPs) and contribute to biological processes such as RNA splicing or transcriptional regulation. These findings suggest that repeat-derived RNAs are important as scaffolds of RBPs and functional elements. However, the overall functional sequences of the repeat-derived RNAs are not fully understood. Here, we show the putative functional repeat-derived RNAs by analyzing the binding patterns of RBPs based on ENCODE eCLIP data. We mapped all eCLIP reads to repeat sequences and observed that 10.75 % and 7.04 % of reads on average were enriched (at least 2-fold over control) in the repeats in K562 and HepG2 cells, respectively. Using these data, we predicted functional RNA elements on the sense and antisense strands of long interspersed element 1 (LINE1) sequences. Furthermore, we found several new sets of RBPs on fragments derived from other transposable element (TE) families. Some of these fragments show specific and stable secondary structures and are found to be inserted into the introns of genes or lncRNAs. These results suggest that the repeat-derived RNA sequences are strong candidates for the functional RNA elements of endogenous noncoding RNAs.

## INTRODUCTION

It is estimated that over half of the human genome consists of repeat sequences, including transposable elements (TEs) (1, 2). In somatic cells, most of these repeat sequences are kept in the transcriptionally repressed state through epigenetic mechanisms (3, 4). On the other hand, fragments of the repeat sequences are transcribed as parts of genes, especially in introns, untranslated regions (UTRs), or long noncoding RNAs (lncRNAs). Recent reports have revealed that repeat-derived RNA sequences embedded in introns or lncRNAs are targets of RBPs and contribute to biological processes such as RNA splicing, localization, or transcriptional regulation(5, 6, 7, 8, 9).

For instance, the LINE1 family, one of the most pervasive TEs that remains actively transposable, is often inserted within the introns of genes in either forward or backward directions (2). Interestingly, fragments derived from the anti-sense strand of young LINE1 sequences can act as scaffolds for the RBP complex, including MATR3 and HNRNPM, which is important for splicing (5). These fragments affect the splicing pattern of the inserted introns. Studies have considered that the insertion of LINEs into the introns of host genes has occurred multiple times during evolution, and as such many subfamilies of LINEs are observed (2, 10). The binding pattern of RBPs is different among the young and old families of LINEs, and thus subsequent effects of splicing also differ among them (5).

LncRNAs are emerging functional RNAs that do not encode proteins, and many of them contain TE sequences (8). In general, lncRNA sequences are less conserved among species compared to coding genes (11, 12), and thus it is difficult to determine the functional region of the sequences. Importantly, at least some lncRNAs seem to have acquired functional domains from TE-derived sequences(7, 8, 9). Xist is a lncRNA of 17 kb in length and plays a pivotal role in X chromosome dosage compensation. Xist has six tandem repeat regions, and all of these regions are thought to derive from TE sequences (13). Recently, it has been shown that one of the tandem repeats, A-repeat, directly binds to Spen, which is an ancient RBP that recruits transcriptional repressors, including histone deacethylases (HDACs) and the NuRD complex (14, 15). Both A-repeat and Spen are required for the X chromosome inactivation (XCI) (16, 17). Importantly, the A-repeat sequence is similar to the fragment of the endogenous retroviral (ERV) K element, which also binds Spen and acts as a transcriptional repressor, suggesting that Xist co-opted TE-derived RNA-protein interactions into the function of lncRNA in XCI (9).

These findings suggest that repeat-derived RNAs are important functional sequence elements; however, these are yet to be fully understood. In general, noncoding RNA functions through its effector proteins and often forms RNA-Protein complexes (18). Thus, it is likely that protein binding sequences in the repeat-derived RNAs are candidates for functional domains of noncoding RNAs, including introns, UTRs, or lncRNAs. To understand which sequences are the targets of an RBP, several UV-crosslinking and immunoprecipitation (CLIP) methods have been developed(19). Among them, enhanced CLIP (eCLIP) was proposed to reveal the precise sequences that bind to an RBP and quantify them(20). To discover the transcriptome-wide protein-binding sequences, over 225 eCLIP experiments were performed in the ENCODE Consortium project(21). These CLIP data are available in the public domain. Utilizing the database, several groups have successfully determined RNA regulatory elements and their binding protein complexes (22, 23). However, frequency of binding of RBPs to repeat-derived RNA sequences were underestimated. This was because raw reads from the sequencing data that map to repeat sequences were discarded in the standardized methods to avoid multi-mapping. To overcome this problem, by analyzing repeat-derived reads of eCLIP, a recent report has been shown that many RBPs actually bind to repeat sequences (24). This analysis focused on the family of repeat sequences, which means representative sequences of TEs, but did not identify either subfamilies or binding positions in the repeat sequences. Importantly, the anti-sense sequence of LINE1 subfamilies shows different RBP binding patterns and functions (5). Therefore, subfamily based and nucleotide-based eCLIP analyses of repeat-derived RNAs are required to identify functional parts of repeat sequences.

Here, we demonstrate the estimation of functional repeat-derived RNAs by analyzing the binding patterns of RBPs based on ENCODE eCLIP data with nucleotide resolution. We have mapped all eCLIP reads to the repeat sequences, including TE subfamilies, and found that multiple RBPs were associated with many repeat sequences. We confirmed the presence of a splicing regulatory complex on the LINE1 anti-sense sequence, which had been reported previously (5) as well as its new candidate for the RBP components. Moreover, we found a new candidate of a functional RNA element that resides at the 3’UTR of LINE1 subfamilies (L1PA4–8), which might be related to the formation of heterochromatin. Furthermore, we found several new RBPs on the fragments derived from other TE families. These fragments show specific and stable secondary structures. Then, we searched for regions in the genome that have these fragments using BLAST (25) search and found that these repeat-derived fragments were inserted into the specific introns of genes or lncRNAs. These results suggest that the repeat-derived RNA sequences are the strong candidates for the functional RNA elements of endogenous noncoding RNAs.

## MATERIALS AND METHODS

### Public eCLIP database and data processing

All eCLIP data were downloaded from the ENCODE Consortium website (https://www.encodeproject.org/) (26). The list of ENCODE accessions for datasets and files are indicated in Supplementary Table 1. Adapters were trimmed from raw reads (cutadapt v2.4)(27) and anything less than 18 bp was discarded following the original protocol(21) with some modifications: Adapter trimming was performed for 3 rounds to remove both 5’ and 3’ adapters. To remove polymerase chain reaction (PCR) duplicates, mapping was first performed against the full human genome (GRCh38) assembly with STAR (v2.6.0c)(28), allowing multi-mapping up to 1,000,000 (–outFilterMultimapNmax 1000000). PCR duplicates were defined as reads (Read1 + Read2), which were mapped on the identical position of the genome and had the same random-mer sequence, with one but all removed. For the multi-mapped reads, PCR duplicates were processed if all of the mapped positions on the genome and its random-mer sequences were identical. The remaining reads (usable reads) were mapped against repeat elements in RepBase (v24.01)(29) with STAR (v2.6.0c). The binding peaks were defined by Piranha (v1.2.1)(30) (bin size = 60 nt) using only Read2 from uniquely mapped reads. Detailed options for each step are indicated in Supplementary Materials and methods. eCLIP data can be recognized as either sense or anti-sense derived signals of repeat sequences using the barcode sequences. The average rate of uniquely mapped/unmapped IP reads to the repeat sequences in Figure 1D were calculated using IP rep1 and IP rep2 data.

**Figure 1.**
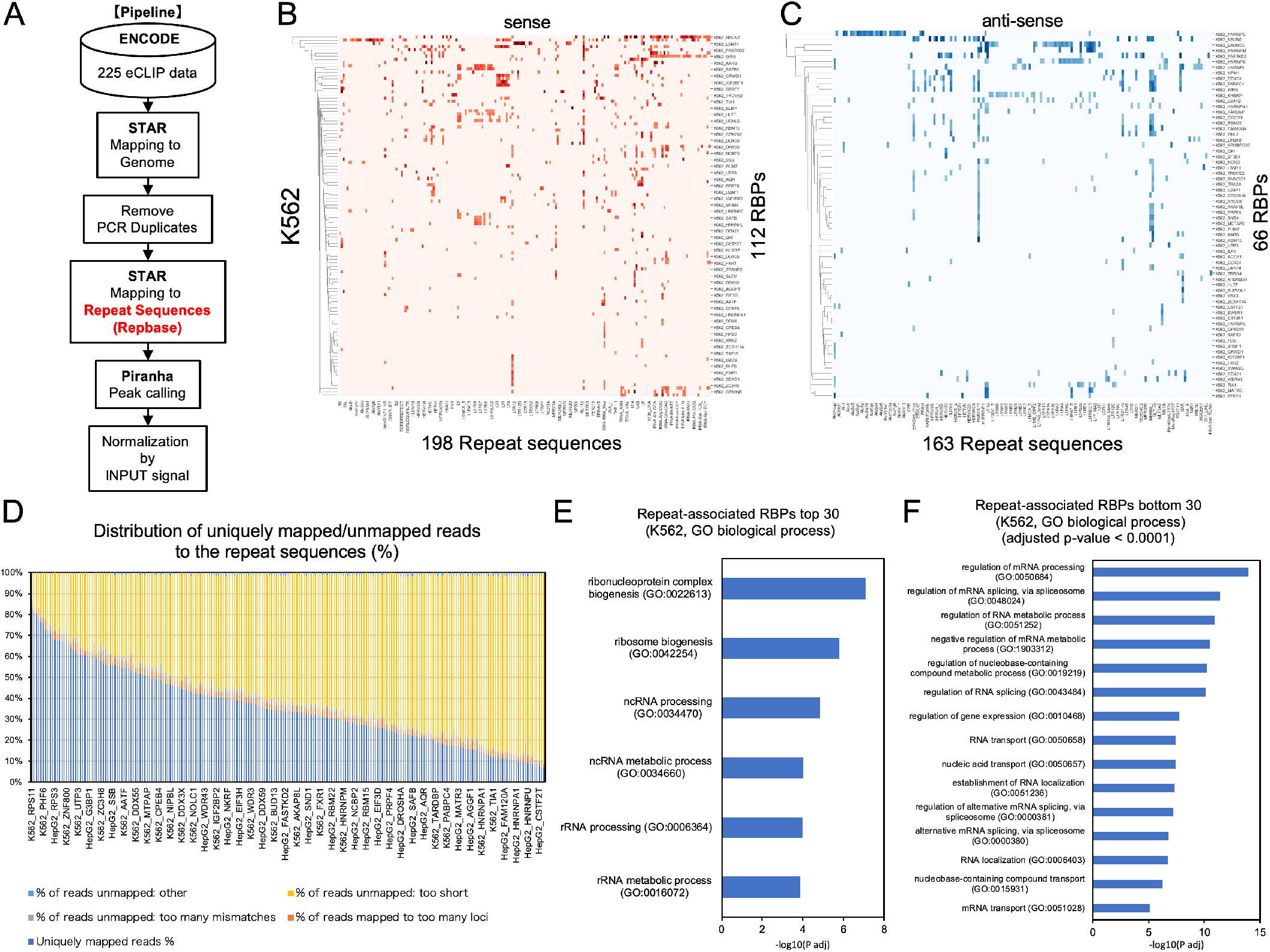
eCLIP data analysis revealed RBP binding patterns to the human RNA repeat sequences. (A) Pipeline of eCLIP data analysis. (B,C) RBP binding pattern of the sense(B) or anti-sense(C) strand of repeat sequences in K562 cells, respectively. The x-axis shows repeat sequences and the y-axis shows RBPs. Red (for sense strand(B)) or blue (for anti-sense strand(C)) colors in the heat maps indicate the maximum values of log2 fold change of eCLIP IP signals compared to SMInput. (D)Distribution of uniquely mapped or unmapped reads to the repeat sequences for 223 eCLIP data (HepG2 or K562). The x-axis indicates the RBPs and the y-axis indicates the percentage of mapped/unmapped reads. (E,F) Gene ontology analysis of the top(E) or bottom(F) 30 of repeat-associated RBPs in K562 cells, respectively. The x-axis shows -log10(adjusted p-value) and the y-axis shows specifically enriched GO terms in the top(E) or bottom(F) 30 of repeat-associated RBPs, which were commonly observed in K562 and HepG2 cells.

### Normalization of eCLIP signal using SMInput

To perform peak-level input normalization, the same pipe line of eCLIP samples was applied to size-matched input controls (SMInput), which are pre-immunoprecipitation samples with an identical size range on the membrane, corresponding to the immunoprecipitation samples (20). SMInput was provided from the original dataset (20, 21). The number of eCLIP reads and corresponding SMInput reads were counted at the nucleotide level and normalized by total usable read counts in each dataset. To reduce data noise, the average read counts for an arbitrary size (every 5 nucleotides) were calculated. The peak-level signal was calculated as log2 fold change (log2[(eCLIP+1)/(SMInput+1)]) of the binding peaks defined by Piranha. For the RBP-binding heat-map data, the threshold was set as over 3 log2 fold-change of eCLIP signals compared to the corresponding SMInput signals. Only the replicate one of two duplicates for each eCLIP sample was used. For the analysis of eCLIP IP read coverage, the peak summit positions were determined using MACS2 (v2.2.6) (31) with the following options (-q 0.01 –nomodel –extsize 30 –call-summits).

### Calculation of RNA secondary structures and its minimum free energy (MFE)

RNA secondary structures were calculated using rtools (http://rtools.cbrc.jp), which includes CentroidFold and CapR (32, 33, 34) with default parameters. CapR calculates the probability of the location of each RNA base within a secondary structural context for a given RNA sequence. Six categories of RNA structures were taken into account: stem part, hairpin loop, bulge loop, internal loop, multi-branch loop, and exterior loop (34). CentroidFold predicts the secondary structure of given RNA sequences and indicates colored bases according to base-pairing probabilities (BPP) and loop probabilities (33). The BPP is a marginal probability indicating that a pair of bases forms a base-pair in a secondary structure. Entire BPPs are efficiently computed using an ensemble of secondary structures by computing the expected probabilities. By using BPPs, we computed the loop probability for a single nucleotide included in a loop structure. In the figure, the warmer colors show higher BPPs and loop probabilities.

The minimum free energy (MFE) for RNA secondary structures was calculated using RNAfold in the ViennaRNA package 2.0 (35). For MFE calculations, RNA fragments were chosen from a part of repeat-derived sequences that consistently formed RNA secondary structures (e.g., hairpin loop) around RBP binding sequences predicted by eCLIP data. For control experiments, identical RNA sequences were randomly shuffled 100 times, preserving dinucleotide counts using uShuffle (36), and their MFEs were calculated using RNAfold. The z-score was calculated as (*m* – *μ*)/*σ*, where *m* is the MFE of a given RNA sequence, *μ* and *σ* are the mean and standard deviations, respectively, of the MFEs of the 100 control sequences. Negative values of z-score show that a sequence is more stable than expected by chance. Prediction of structurally conserved and thermodynamically stable RNA secondary structures among L1PA4-8 was performed using RNAz with the default parameters (37).

### Gene expression data

Gene quantification data in Figure 4G was downloaded from ENCODE RNA-seq data (ENCFF018EJB) (21). Statistical analysis was performed using the Brunner-Munzel test (38).

### BLAST search and Gene ontology analysis

BLAST+(v2.7.1) was downloaded from the NCBI website (ftp://ftp.ncbi.nlm.nih.gov/blast/executables/blast+/LATEST/). The parameter for the length of the initial exact match (word_size) was set to 4. For the analysis of genomic distribution of repeat-derived fragments, human GTF file (GRCh38 release 101) was downloaded from Ensembl website (http://ftp.ensembl.org/). Conversion of GTF to BED file was performed using convert2bed (v2.4.39) (39). Intersection between gene annotation file and BLAST search file was performed using bedtools (v2.29.2) (40). Gene ontology (GO) analysis was performed using gProfiler (the database version: Ensembl 102) (41) or DAVID (v6.8) (42) with default parameters. For gProfiler GO analysis, all human genes were used as the gene background. For DAVID GO analysis, eCLIP 103 RBPs in HepG2 or 120 RBPs in K562 were used as the customized gene backgrounds.

## RESULTS

### eCLIP data analysis revealed RBP binding patterns to the human RNA repeat sequences

To understand which repeat-derived sequences bind to RBPs in human cells, we downloaded ENCODE eCLIP data of HepG2 and K562 cell lines and re-analyzed them. In the original paper of the ENCODE project, repeat-derived sequences were removed before analysis(21). We re-analyzed these repeat-derived sequences following our own pipe line as described below (see Figure 1A). To remove PCR duplicates, we first mapped all pair-end reads of eCLIP to the full human genome (GRCh38) allowing multi-mapping. We defined PCR duplicates as reads that have identical random-mer sequences and genomic positions (5’ to 3’ of Read1 and Read2 region) and removed but one. This is 26.51 % of the all reads. Then, we mapped the remaining reads to the repeat sequence library of Repbase database (v24.01) (29). This library has over 1300 human repeat sequences, including TE subfamilies. To quantify the data, we used only uniquely mapping reads in this step. We confirmed that 35.9 % of the total reads were uniquely mapped to the repeat sequences, whereas 61.32 % on average were unmapped (see Figure 1D and Supplementary table5). We note that only 2.77 % of the total reads were multi-mapped during this step because this library has fewer redundant sequences (see Figure 1D). Consistent with the previous report, this result suggests that a large part of the reads is derived from repeat sequences and that many RBPs are binding to the repeat elements(24). We also checked the fraction of SMInput reads of each RBP mapped to the repeat elements (see Supplementary Figure 1A,B and Supplementary table5). We found that 49.0% on average of the SMInput was mapped to repeat elements. This result suggests that many RBPs actually bind to repeat sequences because the SMInput is not an empty control, but it contains many RBP-binding fragments (RBP footprint) in addition to the target IP RBPs. Then, we defined RBP binding peaks using a peak caller, Piranha(30). To quantify the mapping signals of binding peaks, we processed SMInput data identically to the eCLIP data and calculated the relative read number for each position (see Materials and Methods). To reduce the noise in the data, we conducted data smoothing using a bin for every 5 nucleotides.

To investigate what fraction of the original reads actually represented binding events to the repeat sequences, we calculated the fraction of reads that were mapped to the repeat sequences and enriched compared to SMInput reads. We defined the IP reads in the peak-called peaks, which were enriched at least over SMInput reads, contributing to the binding events in each RBP. Then, we calculated IP read fractions that were over 1-, 2-, 4-, and 8-fold over SMInput reads using 5 nt bins of IP rep1 read coverage data (see Supplementary Figure 1C,D,E). We found that 20.95 %, 10.75 %, 4.47 %, and 1.86 % on average of the IP reads were at least over 1-, 2-, 4-, and 8-fold over SMInput in K562 cells, respectively. Similarly, we found that 15.35 %, 7.04 %, 2.69 %, and 1.37 % on average of the IP reads were at least over 1-, 2-, 4-, and 8-fold over SMInput in HepG2 cells, respectively. The percentage of substantial binding reads varied among RBPs, ranging from approximately 0% to 70%. These results suggest that a considerable population of the total IP reads contributes to the binding events on the repeat sequences.

To understand which RBPs are associated with the repeat sequences, we performed gene ontology (GO) analysis for the top or bottom 30 RBPs, which showed a higher or lower percentage of substantial binding reads to the repeat sequences, respectively, using gProfiler (41) (see Figure 1E, F Supplementary Figure1F-I, Supplementary Table2). We observed that the top 30 genes were explicitly enriched with GO terms, including ‘rRNA processing (GO:0006364)’ or ‘ncRNA processing (GO:0034470)’, which were not observed in the bottom 30 genes for both HepG2 and K562 cells. This is because the repeat sequence library we used contains many kinds of rRNAs as well as other ncRNAs including snRNA, snoRNA, and tRNA, and the top 30 RBPs contain ribosomal proteins. Conversely, the bottom 30 genes were specifically enriched with GO terms, including ‘RNA transport (GO:0050658)’ and ’RNA localization (GO:0006403)’, which were not observed in the top 30 genes for both HepG2 and K562 cells. (see Figure 1F and Supplementary Figure1F,G,I, Supplementary Table2). To further investigate the characteristics of the top or bottom 30 RBPs, we also analyzed enriched GO terms of these RBPs considering all eCLIP RBPs in each cell line as background using another GO analysis tool, DAVID. We obtained consistent results that show ‘rRNA processing (GO:0006364)’ or ‘mRNA processing (GO:0006397)’ was significantly associated with the top or bottom 30 RBPs, respectively, over the background in both HepG2 and K562 cells (see Supplementary table2). These results suggest that the binding tendency to the repeat sequences was the property of RBPs, which can relate to their functions.

To estimate how many RBPs were significantly binding to the repeat sequences, we set the threshold value at over 3 of the log2 fold change to consider strong signals only. We then counted the number of repeat sequences (sense strand) that have at least one RBP binding signal. We found that 149 repeat sequences bind to 87 RBPs in HepG2 cells and 198 repeat sequences bind to 112 RBPs in K562 cells (see Figure 1B and Supplementary Figure1J,L). We also searched for the anti-sense of repeat sequences that bind to RBPs and found that 132 repeat sequences bind to 48 RBPs in HepG2 cells and 163 repeat sequences bind to 66 RBPs in K562 cells (see Figure 1C and Supplementary Figure1K,M).

### LINE subfamilies show specific and different binding patterns of RBPs

Next, we focused on RBP binding patterns of the repeat sequences on each TE subfamily. Using eCLIP data, we visualized binding patterns of each TE subfamily to RBPs as heat-maps (see Supplementary Figure2). The LINE family contains several subfamilies, including L1P (primate-specific) and L1M (mammalian-wide). L1PA1 (including L1 and L1HS in Repbase), a sub-class of L1P, is the youngest and only active LINE subfamily in the human genome. The binding heat-map of LINE sense strands showed that most of the RBPs bind to the younger subfamily L1P(see Figure 2A). Consistent with the previous report of the ENCODE project(24), we confirmed that L1 binds to L1 suppressor proteins including SAFB, PPIL4, and TRA2A in K562 cells. In addition to that, we confirmed that sense strands of L1 and L1Ps also bind to several L1 activators including SLTM, SRSF1, and suppressor HNRNPL in both HepG2 and K562 cells (see Figure 2A, C)(43, 44). This result suggests that our method is reliable and significantly recaptures true positive regulatory RBPs that bind to L1 in the cell. Interestingly, a series of subfamilies (L1PA4-L1PA8) showed similar and specific RBP binding patterns in both HepG2 and K562 cells, suggesting that subfamilies of L1 might have different binding partners and functions (see Figure 2A)(we will discuss this further later).

**Figure 2.**
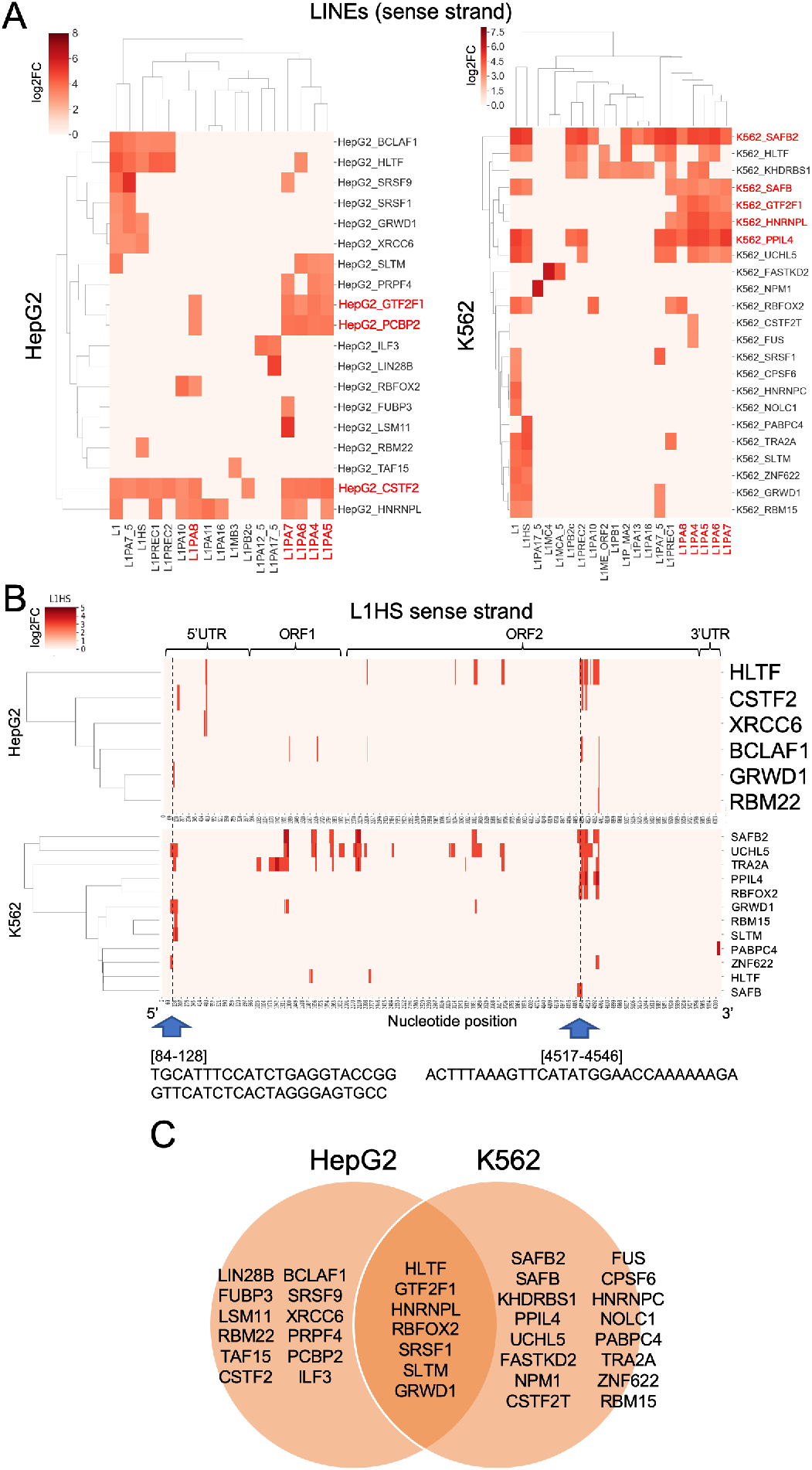
LINE subfamilies show specific and different binding patterns of RBPs. (A)RBP binding pattern on LINE subfamilies. The x-axis indicates the sense strand of the LINE subfamilies and the y-axis indicates the RBPs. Red colors indicate the maximum values of log2 fold change of eCLIP IP signals compared to SMInput. RBPs written in red letters indicate common binding RBPs among L1PA4-8 in each cell. (B)RBP binding pattern on the L1HS sense strand. The x-axis indicates the nucleotide positions of L1HS and the y-axis indicates binding RBPs. Red colors indicate log2 fold change of eCLIP IP signals compared to SMInput. (C)Venn diagram for RBP binding to LINE subfamilies.

### Analysis of RBP binding patterns on young active LINE1 sequences with nucleotide resolution revealed short stable RNA fragments with multiple binding of RBPs

To understand which sequence is important for the LINE1 regulation, we investigated the RBP binding patterns on the L1HS sequence with nucleotide resolution. We calculated and visualized the binding signals of every 5 nucleotides on the L1HS sequence (see Figure 2B). On the L1HS sense strand, we found that several ‘hot spot’ sites with multiple RBPs could bind together and/or separately. We hypothesized that these sites are local binding sites of RBP clusters and potential RNA regulatory sequences.

To investigate which sequences actually bind to RBPs using nucleotide resolution, we first checked eCLIP IP read coverage of clustered RBP binding sites on the L1HS 5’UTR (see Figure 2B and 3A). The 5’ end of the read2 sequence of eCLIP is considered the binding point between the RNA and protein (20). Therefore, we focused on short sequences from the binding peak summit to the 5’ upstream end of the binding peak, as indicated by the red rectangle in Figure 3A. We found that this region contains a putative binding sequence (CAUCUGAG) of GRWD1 and ZNF622, which is similar to the common consensus motif (CAUCGAG) (23). Then, we investigated whether this region forms specific RNA secondary structures. We checked RNA secondary structures both upstream and downstream of the peak summit average [position 132] and found that position 84-128 (termed L1HS[84-128]) was consistently predicted to form a stable hairpin loop structure (see Supplementary Figure 3A-D and Figure 3B). Notably, GRWD1 and ZNF622 were predicted to bind to the bulge-containing stem sequence. Similarly, we focused on another RBP-binding hot spot on the L1HS ORF2 (see Figure 2B) and found that the 5’ region of the clustered RBP-binding peaks contains putative binding sequences of SAFB (GGAAAAA), UCHL5 (AUGGAACC), and PPIL4 (GAGCCCG) that were similar to each of the consensus motifs (SAFB:GGAAAGA, UCHL5:AUGGACC, PPIL4:AAGCCCG)(see Figure 3E)(23). We also checked the RNA secondary structure of this region and found that position 4517-4546 on L1HS (termed L1HS[4517-4546]) was predicted to form a stable hairpin loop structure (see Supplementary Figure3E-H and Figure 3F). UCHL5 was predicted to bind to the hairpin and stem region, whereas SAFB2 and PPIL4 seemed to be associated with the flanking region of the hairpin loop structure (see Figure 3E,F). To investigate whether these fragments were significantly stable, we calculated the minimum free energy (MFE) using RNAfold and compared them to randomly shuffled sequences with preserved dinucleotides of each fragment (see Figure 3C,G). We found that the MFE of L1HS[84-128] (−15.10 kcal/mol) was significantly lower than the average MFE of 100 shuffled sequences (−9.222 kcal/mol)(z-score=-2.555). Similarly, the MFE of L1HS[4517-4546])(−2.80 kcal/mol) was significantly lower compared to the average MFE of 100 shuffled sequences (−2.138 kcal/mol)(z-score=-0.368). These results suggest that these fragments can form stable structures.

**Figure 3.**
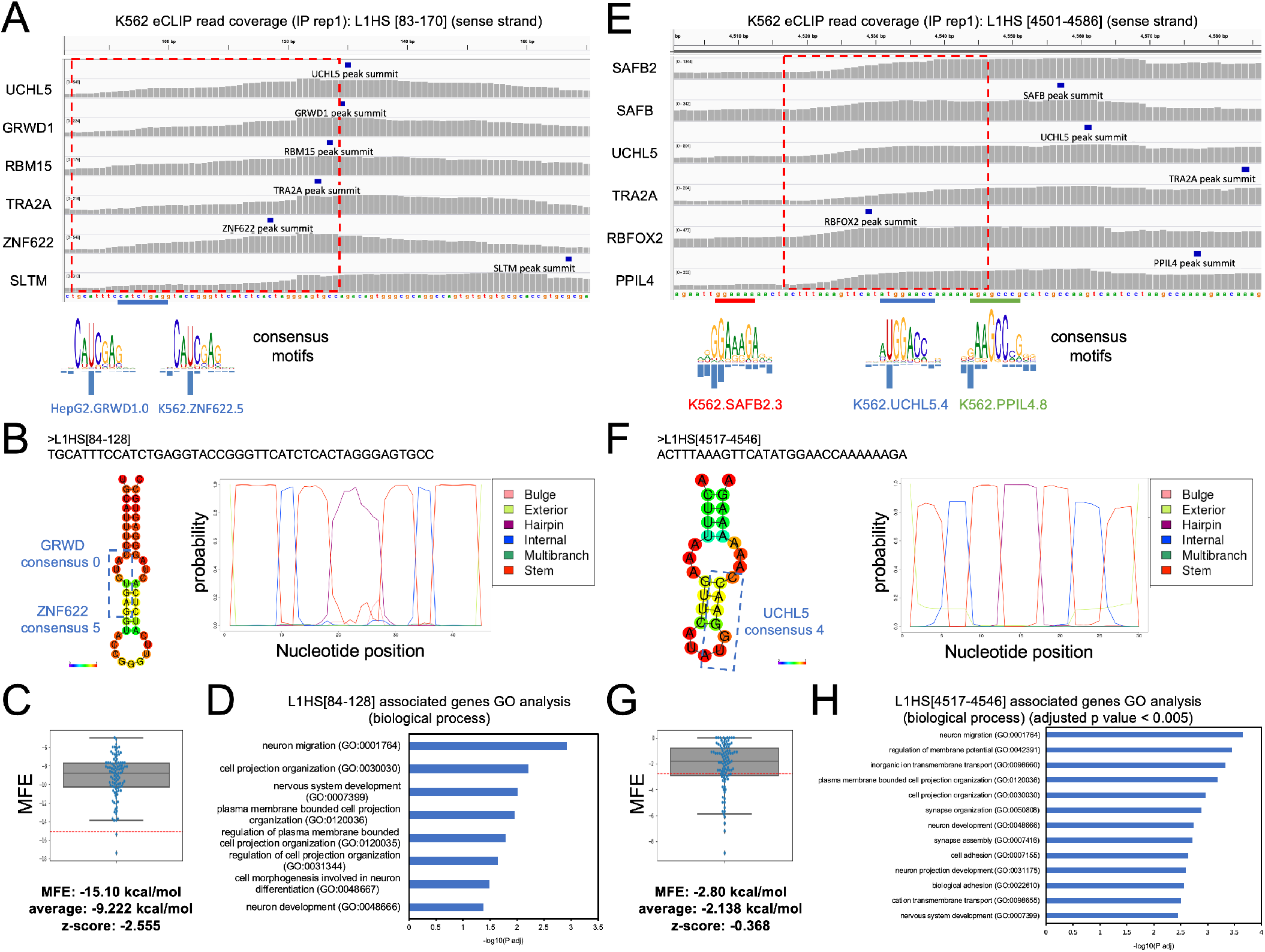
Analysis of RBP binding patterns on L1HS with nucleotide resolution revealed short stable RNA fragments with multiple binding of RBPs (A,E)eCLIP IP read coverage of RBPs that are associated with L1HS[84-128](A) or L1HS[4517-4546](E) in K562 cells. The y-axis shows the RBP names, and the x-axis shows the nucleotide position of the L1HS sequence. The red rectangles indicate the RBP binding sites of L1HS[84-128](A) or L1HS[4517-4546](E), which were predicted to form stable RNA secondary structures shown in (B,F), respectively. Putative RBP-binding sites of GRWD1, ZNF622(A), SAFB, UCHL5, and PPIL4(E) are highlighted with the blue(A), red, blue, and green(E) bars, respectively. RBP consensus motifs were adapted from mCrossBase (23) and compared with putative RBP binding sites. (B,F)Prediction of RNA secondary structures for RBP cluster sites of L1HS[84-128](B) or L1HS[4517-4546](F) calculated using CentroidFold (left panels) or CapR (right panels). The colored bases in the left panels (CentrioidFold) indicate base-pairing and loop probabilities. The blue rectangle shows putative RBP binding sites. The right panel (CapR) shows the structural profile of an RNA base by a set of six probabilities (stem part, hairpin loop, bulge loop, internal loop, multibranch loop, and exterior loop) that the base belongs to each category. The x-axis indicates the nucleotide position and the y-axis indicates the probability of the structural profiles. (C,G)Distribution of MFE for randomly shuffled sequences of L1HS[84-128](C) or L1HS[4517-4546](G). The y-axis indicates MFE (kcal/mol). Horizontal red bars indicate MFE of L1HS[84-128](C) or L1HS[4517-4546](G), respectively. The z-score and average MFEs were calculated as shown in the Materials and Methods section. (D,H)Gene ontology analysis of L1HS[84-128](D) or L1HS[4517-4546](H) inserted genes. The x-axis indicates -log10 (adjusted p-value).

Next, we inquired which transcripts were the origins of these fragments. Because eCLIP reads are too short due to the technical process, it is very difficult to reconstruct the original RBP binding sequences from the read data, especially in repeat-containing regions. There are two possibilities: these fragments are derived from active L1HS transcripts and/or transcripts that contain L1HS fragment sequences. Although L1HS binding RBPs include several known regulators of L1HS, it is still possible that parts of the eCLIP signals were derived from the fragments of L1-containing genes. Therefore, we searched for genes that contain consensus sequences of the L1HS[84-128] or L1HS[4517-4546] using BLAST to predict potential original sequences. We obtained 268 and 1298 candidate genes containing each L1HS fragment, respectively. Most of these fragments were inserted into introns of genes (see Supplementary table3). To investigate the association of genes with L1HS[84-128] or L1HS[4517-4546], we performed GO analysis of these transcripts. We found that L1HS[84-128]-associated genes were significantly enriched in specific GO terms, such as ‘neuron migration (GO:0001764)’, ‘cell projection organization (GO:0030030)’, and ‘nervous system development (GO:0007399)’ (see Figure 3D and Supplementary table4). Similarly, L1HS[4517-4546]-associated genes were also significantly enriched in specific GO terms such as ‘regulation of membrane potential (GO:0042391)’, and ‘synapse organization (GO:0050808)’ (see Figure 3H and Supplementary table4). These results suggest that L1HS[84-128] and L1HS[4517-4546] are inserted into specific genes that regulate neurogenic functions.

Interestingly, it has been shown that the position from around 80-100 of the L1 sense strand, which overlaps with L1HS[84-128], was the binding site for transcription factors and regulated L1 transcription(45). Together, these results support the notion that RNA fragments of RBP binding hot spots are strong candidates for RNA regulatory elements.

### Analysis of RBP binding patterns on L1PA subfamilies with nucleotide resolution revealed putative functional RNA elements that associate with the RBP complex

To investigate the difference of RBP binding patterns among older LINE1 subfamilies, especially L1PAs, we visualized RBP binding patterns on L1PA subfamilies (see Figure 2A and 4A, Supplementary Figure4A). Interestingly, L1PA4-8 subfamilies specifically bind to a common set of RBPs on specific regions around position 750 to 800, which is included within the 3’UTR region of LINE1 subfamilies in both HepG2 and K562 cells. This complex contains CSTF2, PCBP2, GTF2F1 in HepG2 cells and GTF2F1, SAFB, SAFB2, PPIL4, HNRNPL in K562 cells. Notably, this complex is very similar to the RBPs that bind to 4517-4546 of the L1HS sense strand in K562 cells but uniquely contains GTF2F1 (see Figure 2B), suggesting that this complex might be a part of the functional unit.

**Figure 4.**
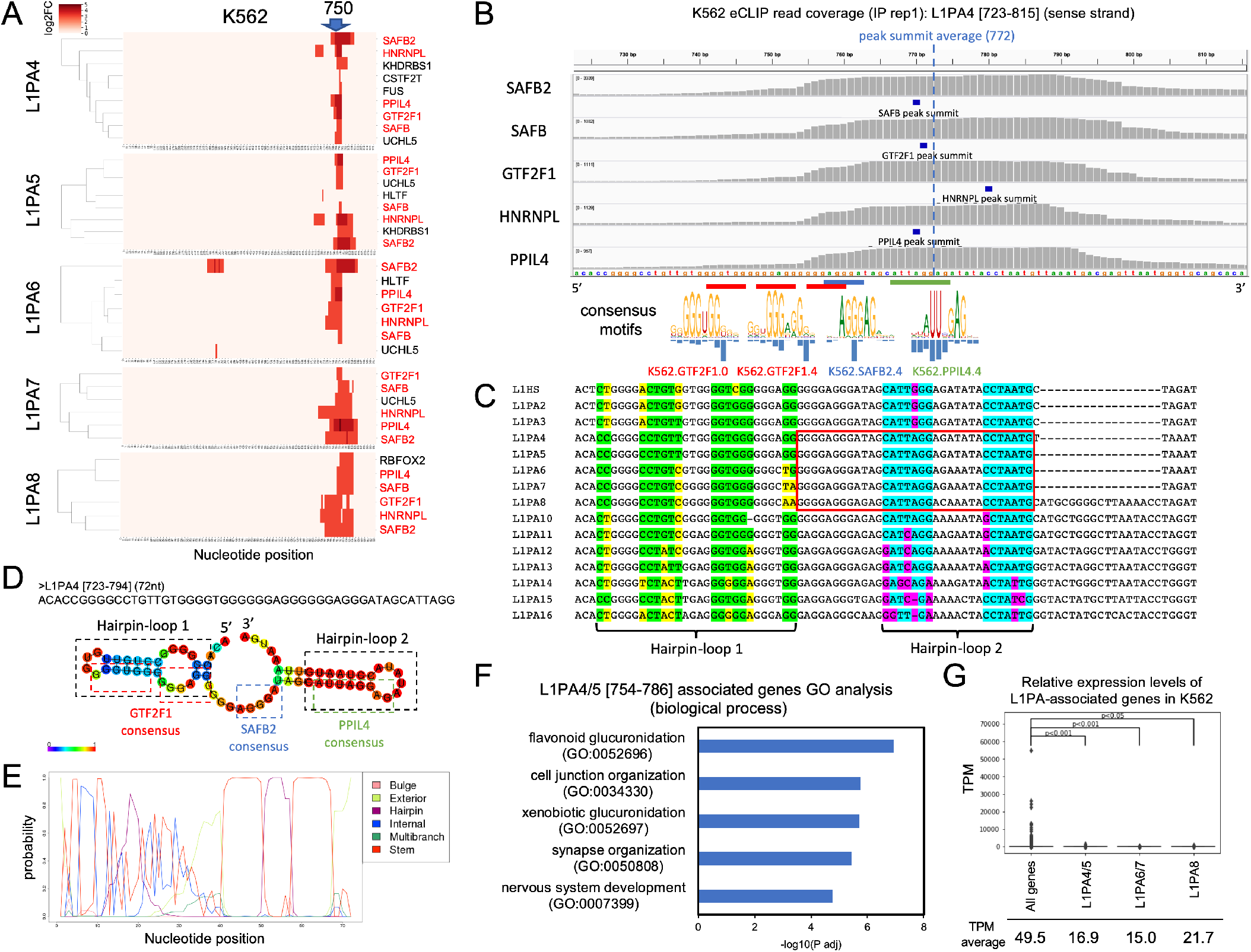
Analysis of RBP binding patterns on L1PA subfamilies with nucleotide resolution revealed putative functional RNA elements that associate with the RBP complex.(A)RBP binding pattern of L1PA4-8 sense strand (3’ fragments) with nucleotide resolution in K562 cell. The x-axis indicates the nucleotide position of each RNA. Red colors in the heat maps indicate a log2 fold change of eCLIP IP signals compared to SMInput. RBPs written in red letters indicate common binding RBPs among L1PA4-8 in K562 cell. (B)eCLIP IP read coverage of RBPs that associate with L1PA4 in K562 cells. The y-axis shows the names of RBPs, and the x-axis shows the nucleotide position of the L1PA4 sequence. The blue vertical bar indicates the average position of the RBP binding peak summit. Putative RBP-binding sites of GTF2F1, SAFB2, and PPIL4 are highlighted with the red, blue, and green bars, respectively. RBP consensus motifs were adapted from mCrossBase (23) and compared with putative RBP binding sites. (C)Multiple sequence alignments of L1PA subfamilies associated with the RBP complex. The green and sky blue letters indicate conserved nucleotides between L1PA4 and L1PA5 that were predicted to form base pairing for hairpin-loop structures, whereas the yellow and magenta letters indicate mutations in other L1PA subfamilies for hairpin-loop structures. The red rectangle indicates RNA fragments that were used to search for associated genes in (F) and (G). (D,E) RNA secondary structure prediction of L1PA4[723-794] calculated using CentroidFold(D) or CapR(E). (D) Putative RBP-binding sequences of GTF2F1 (red), SAFB2 (blue), and PPIL4 (green) or hairpin loop structures (black) are indicated by colored rectangles. Colored bases indicate base-pairing and loop probabilities. (F) Gene ontology analysis of genes associated with L1PA4[754-786]. The x-axis indicates the -log10(adjusted p-value). (G)Relative gene expression levels of L1PA-associated genes in K562 cells. The y-axis indicates transcripts per million (TPM). The average TPM of L1PA-associated genes are indicated below. P-values of the Brunner-Munzel test between each sample are indicated (38).

To search for the RBP binding candidate sequences in nucleotide resolution, we checked eCLIP IP read coverage of RBPs associated with L1PA4 in K562 cell. We found multiple binding candidate sequences of GTF2F1 (GGGUGG, GGGAGG; consensus motif: GGGUGG, GGGAGG), SAFB2 (AGGGAU; consensus motif: AGGGAG), and PPIL4 (AUUAGGAG; consensus motif: AUUNGAG), all of which were very similar to each binding consensus motif, in the 5’ upstream region of the RBP binding peak summit on L1PA4 (see Figure 4B)(23). We then asked whether this region forms RNA secondary structures and found that this region was predicted to form two hairpin loop structures: hairpin loop 1 is formed on the 5’ upstream region of the binding peaks of RBPs and hairpin loop 2 overlaps with the same peaks (see Figure 4C,D,E and Supplementary Figure4B,C). It appears that GTF2F1 binds to the hairpin and stem or bulge region of the hairpin loop 1, and SAFB2 binds to the flanking region between hairpin loops 1 and 2, whereas PPIL4 binds to the hairpin and stem region of hairpin loop 2 (see Figure 4B,C,D). In addition, hairpin loop 2 seems to be more stable than hairpin loop 1 (see Figure 4D,E).

Because L1PA4 binding RBPs are also associated with L1PA5-8, we checked the sequence conservation among L1PA subfamilies of the corresponding regions (see Figure 4C). We noticed that all binding candidate sequences of RBPs were conserved among L1PA4-8 but also other L1PA subfamilies, including L1PA2 and L1PA3. Interestingly, we noticed that the stem region of the hairpin loop 2 (indicated by sky blue letters in Figure 4C) was completely conserved among the L1PA4-8 subfamily, whereas that of hairpin-loop 1 (indicated by green letters in Figure 4C) harboured several mutations (indicated as yellow letters) among the L1PA subfamilies. To determine whether these L1PA4-8 regions form conserved and thermodynamically stable RNA secondary structures, we investigated each of the sequences and compared them using Centroidfold, CapR, and RNAz (see Supplementary Figure4D,E) (37). We found that hairpin loop 2 was stable and conserved among the L1PA4-8 subfamily, but hairpin-loop 1 was relatively unstable and did not consistently form secondary structures. We also checked the RNA secondary structure of other L1PA subfamilies, including L1PA3 and L1PA10 (see Supplementary Figure4F), and found that these sequences could also form similar secondary structures, although they seemed relatively unstable. These results suggest that in addition to the RBP binding consensus sequence, RNA secondary structures and their thermodynamic stability, which could interfere with surrounding sequences and structures, are essential for RBP binding and RNA-RBP complex formation.

Next, to understand whether these L1PA4-8 fragments could act as functional elements for gene regulation, we first searched for genes associated with these elements using BLAST. We identified a potentially essential sequence (L1PA4-8[754-786]), which contains RBP binding motifs and a conserved hairpin loop 2 structure as a putative functional element. We searched a 100% match sequence against the human genome and obtained 1358, 617, and 175 candidate genes associated with L1PA4/5, L1PA6/7, and L1PA8, respectively (see Supplementary table3). Most elements were inserted into the introns of the genes. GO analysis revealed that these elements were significantly associated with specific GO terms related to neuron and synapse organization, suggesting that L1PA fragments were preferentially inserted and retained in a set of functionally related genes (see Figure 4F and Supplementary table4).

To investigate the effect of these potential functional elements on gene regulation, we checked the expression levels of these L1PA-associated genes using ENCODE RNA-seq data. We found that the expression levels of all L1PA4-8 associated genes were significantly lower than the average of all gene expression levels (p<0.05, Brunner-Munzel test (38)) (see Figure 4G). This result is consistent with a previous report showing that the insertion of the LINE1 element into introns of genes inhibits host gene expression levels (43). Furthermore, these elements bind to RBPs, including SAFB, SAFB2, and HNRNPL, which have been reported to repress gene expression and are associated with heterochromatic regions (43, 46, 47, 48, 49, 50). These results suggest that these RBPs may recognize specific RNA structures to form an RNA-protein complex on the L1PA fragments, resulting in the formation of heterochromatic regions and low expression levels of the inserted genes. Taken together, we conclude that the sense strand of L1PA4-8 may have some RNA sequence modules that act as scaffolds for RBP binding and complex formation, which act as a repressive RNA element.

### Anti-sense of LINE1 subfamilies shows distinct binding pattern of RBPs from sense strand LINEs

Previous reports have also revealed that distinct RBPs from the L1 sense strand bind to the L1 anti-sense strand and have function of splicing functions(5, 24). We also confirmed that the components of a large assembly of splicing regulators (LASR), HNRNPM and MATR3 bind to the L1 anti-sense sequence(51)(see Figure 5A and Supplementary Figure5A). We also confirmed that the stress granule factor TIA1 and splicing factor PTBP1 bound to the L1 anti-sense sequence. PTBP1 was reported as a co-regulator of MATR3(52). Interestingly, TIA1 was also reported as a co-regulator of PTBP1, suggesting that these proteins are associated with the L1 anti-sense sequence together and regulate splicing mechanism(53).

**Figure 5.**
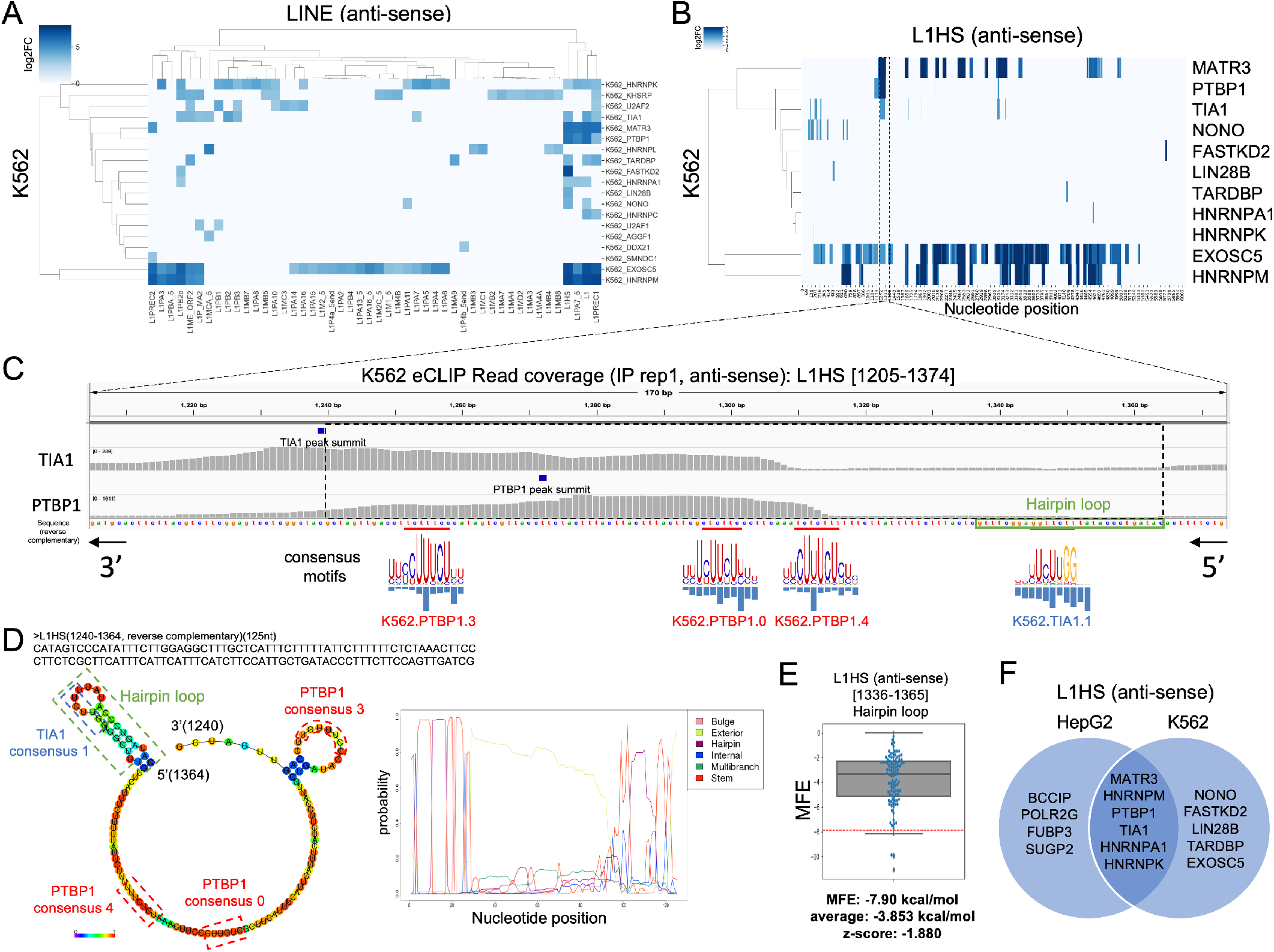
Anti-sense of LINE1 subfamilies shows distinct binding pattern of RBPs from sense strand LINEs (A)RBP binding pattern on anti-sense of LINE1 subfamilies in K562 cells. The x-axis indicates the LINE subfamilies (anti-sense strand) and the y-axis indicates RBPs. Blue colors indicate the maximum values of log2 fold change of eCLIP IP signals compared to SMInput. (B)RBP binding pattern on anti-sense of L1HS with nucleotide resolution in K562 cells. The x-axis indicates the nucleotide position of the anti-sense of L1HS and y-axis indicates RBPs. Blue colors indicate log2 fold change of eCLIP IP signals compared to SMInput. (C)eCLIP IP read coverage of RBPs that are associated with L1HS[1205-1374](anti-sense) in K562 cells. The y-axis shows the names of RBPs, and the x-axis shows the nucleotide position of the L1HS sequence. The black rectangle indicates the putative RBP binding region that was used to predict RNA secondary structures in (D). The green rectangles in (C, D) indicate the sequence of a hairpin loop structure that overlap with L1HS-AS[1336-1365]. Putative RBP binding sites of PTBP1 and TIA1 are highlighted with red and blue bars, respectively. RBP consensus motifs were adapted from mCrossBase (23) and compared with putative RBP binding sites. (D)Prediction of RNA secondary structures for RBP binding sites of anti-sense L1HS[1240-1364] calculated by CentroidFold(left panel) or CapR(right panel). The colored bases in the left panel indicate the base-pairing and loop probabilities. The x-axis of the right panel indicates the nucleotide position of anti-sense L1HS[1240-1364], and the y-axis indicates the probability of the structural profiles.(E)Distribution of MFE for 100 randomly shuffled sequences of anti-sense L1HS[1336-1365] that forms a hairpin-loop structure shown in green rectangles in (C, D). The y-axis indicates MFE(kcal/mol). Horizontal red bar indicates MFE of anti-sense of L1HS[1336-1365]. The z-score and average MFEs were calculated as shown in the Materials and Methods section. (F)Venn diagram for RBP binding to anti-sense L1HS in HepG2 and K562 cell.

To understand the precise mechanism of these RBP binding, we investigated the RBP binding pattern to the L1HS anti-sense sequence with nucleotide resolution. Consistent with previous reports(24), MATR3 and HNRNPM showed strong signals over a wide range of the L1HS anti-sense sequence (see Figure 5B and Supplementary Figure5B). SUGP2 and EXOSC5 also showed very similar binding patterns to MATR3 and HNRNPM, suggesting that these proteins are also integral components of this complex. In contrast to that, PTBP1 and TIA1 showed narrow binding peaks. This might reflect the different binding modes of these RBPs, namely, broad binding and specific binding, and thus different mechanisms or functions in the complex. We then focused on the binding sites of PTBP1 and TIA1 and investigated the characteristic features of this fragment. eCLIP IP read data revealed that the PTBP1 peak region contains at least three binding candidate sequences of PTBP1 (UUUUCUCU, CCUUCUC, and CCUUUCU), which were very similar to the consensus binding motifs (CUUUCUCU, UCUUCUC, and CCUUUCU) (see Figure 5C)(23). Moreover, the 5’ upstream region of the TIA1 and PTBP1 peaks contains a TIA1 binding consensus motif (UUCUUGG)(23). Then, we estimated the RNA secondary structures of this RBP binding region (L1HS anti-sense position [1240-1364]) using CentroidFold and CapR and found that the TIA1 binding fragment can clearly form a hairpin-loop structure (termed L1HS-AS[1336-1365], indicated by green rectangles in Figure 5C,D) (see Figure 5C,D and Supplementary Figure5C-G). We calculated the MFE of L1HS-AS[1336-1365] and compared it with those of randomly shuffled sequences of the fragment; we found that L1HS-AS[1336-1365] can form a stable structure (MFE= −7.90 kcal/mol, z-score= −1.880) (see Figure 5E). In contrast, the PTBP1 binding candidate sequences were not predicted to form strong RNA secondary structures (see Figure 5C,D). These results suggest that both secondary structures and sequence motifs are important for RBP binding, and multiple tandem binding sequences of RBPs are important for forming an RNA-protein complex.

In addition to the above RBPs, we also observed that BCCIP, POLR2G, FUBP3 in HepG2 cells, NONO, FASTKD2, LIN28B, TARDBP in K562 cells, and HNRNPA1 and HNRNPK in both cellsbind to the L1HS anti-sense sequences (see Figure 5B,F and Supplementary Figure5B). These RBPs might be additional regulators of the sequence. Taken together, these results strongly support a previously proposed notion that fragments derived from the L1 anti-sense sequence act as a scaffold for the regulatory RBP complex, and this region contains a candidate for RNA functional elements.

### Analysis of RBP binding patterns on sense or anti-sense of HERV subfamilies with nucleotide resolution predicts putative functional RNA elements

We also analyzed major TE subfamilies, including ALU, HERV, MER, LTR, and SVA (see Supplementary Figure2). Interestingly, these repeat sequences showed specific RBP binding patterns among subfamilies and strands. For instance, the binding heat-map of HERV sense or anti-sense strands showed that the binding patterns of RBPs are different among HERV subfamilies (see Figure 6A, C 7A, F). To investigate putative functional RNA fragments, we focused on the sequences that showed multiple binding signals on the sense strand of HERV39(see Figure 6B). Interestingly, IGF2BP1 and SRSF7 specifically bind to the simple repeat sequence (..UGUGUGUGU..) in the HERV39 sense strand in both HepG2 and K562 cells, which is consistent with one of the SRSF7 binding consensus motifs (GUGUGUG)(23). In K562 cells, we observed that this UG repeat sequence is also associated with TARDBP and EIF4G2, which have the same consensus motif (GUGUGUG)(23). Notably, it has been reported that TARDBP preferentially binds to UG repeat sequences (54, 55, 56) and is associated with cryptic exons, as well as SRSF7, TIA1, and IGF2BP1 (57, 58). Other groups have reported that TARDBP could bind to IGF2BP1 and SRSF7 (59, 60, 61), suggesting that these RBPs may cooperatively or competitively regulate cryptic exons through UG repeat sequences. To estimate which sequences are the origin of this fragment, we searched for similar transcriptome fragment sequences (bit score over 80) using BLAST. We obtained 1084 HERV39[3907-3962] associated gene candidates and found that most of these fragments reside in introns of genes, including MAML2 (see Figure 6D and Supplementary Table3). These data suggest that this fragment might have some function during splicing regulation and RNA stabilization. GO analysis revealed that neuron and synapse related GO terms including ‘neuron differentiation (GO:0030182)’ were highly enriched with these genes (see Supplementary Figure6A,B and Supplementary Table4), which is consistent with previous reports (62, 63).Similarly, we found that RBP binds to a hot spot in the anti-sense strand of HERVS71 (see Figure 7B). We predicted the RNA secondary structure of this fragment and found that this sequence also forms a hairpin loop structure (see Figure 7C, D and Supplementary Figure7A-C). We found one example that this fragment was inserted into an intron of the DET1 gene (see Figure 7E).Importantly, this fragment binds to the miRNA processing complex, including DROSHA and DGCR8, suggesting that this fragment could be a pre-miRNA. Overall, these results show that our approach is promising in search of candidates for repeat-derived functional RNA elements.

**Figure 6.**
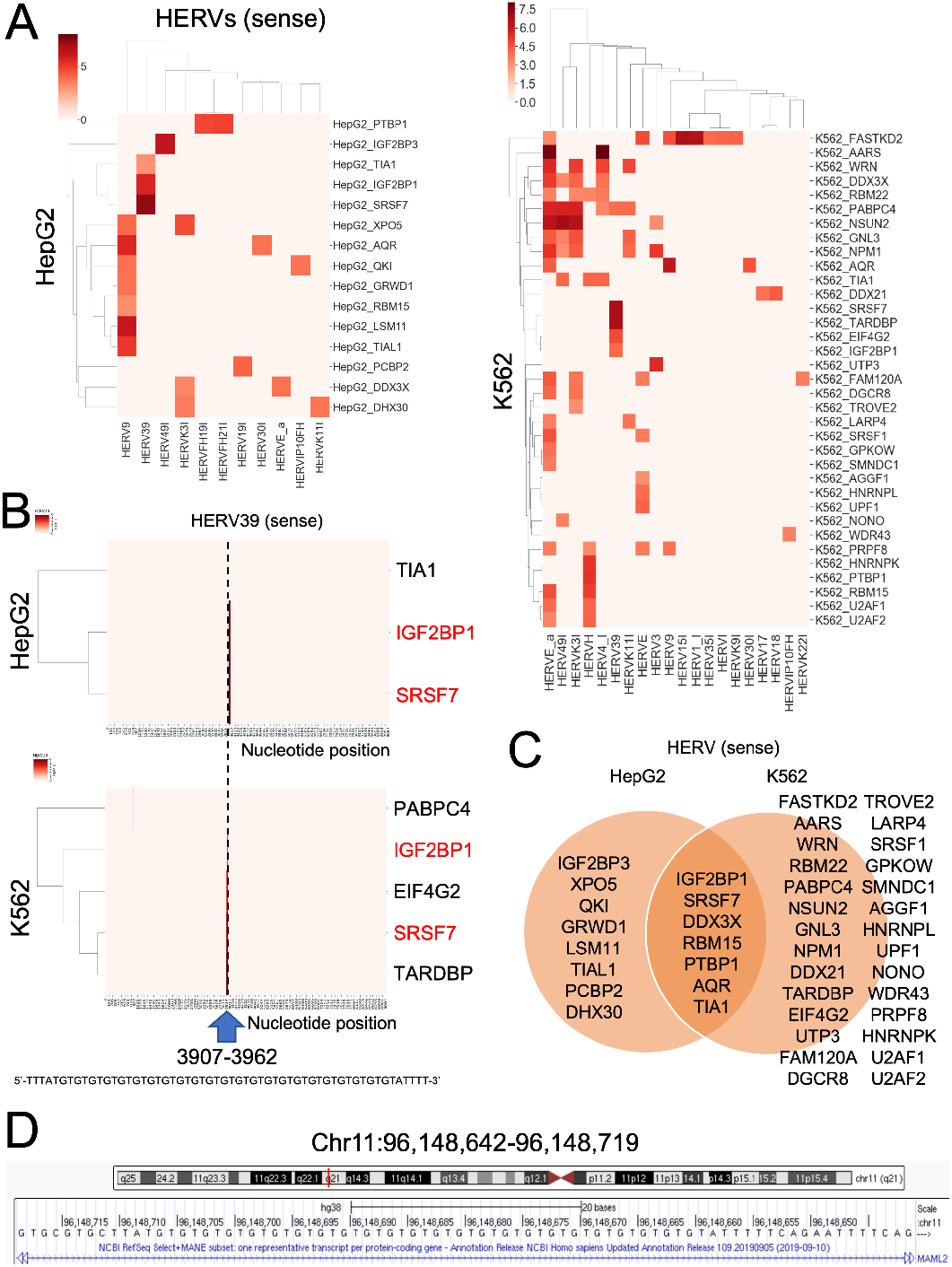
Analysis of RBP binding patterns on HERV subfamilies with nucleotides resolution predicts putative functional RNA elements. (A)RBP binding pattern of the sense strand of HERV subfamilies. The x-axis indicates the sense strand of HERV subfamilies and the y-axis indicates RBPs. Red colors indicate the maximum values of log2 fold change of eCLIP IP signals compared to SMInput.(B)RBP binding pattern of HERV39 sense strand with nucleotide resolution. The x-axis indicates the nucleotide position of HERV39 and the y-axis indicates RBPs. Red colors indicate log2 fold change of eCLIP IP signals compared to SMInput. RBPs written in red letters indicate common binding RBPs between HepG2 and K562. (C)Venn diagram for RBP binding to HERV subfamilies. (D)Example of HERV39(3907-3962) in the intron of gene. HERV39(3907-3962) sequence in the first intron of MAML2 gene is indicated in the UCSC genome browser (hg38).

**Figure 7.**
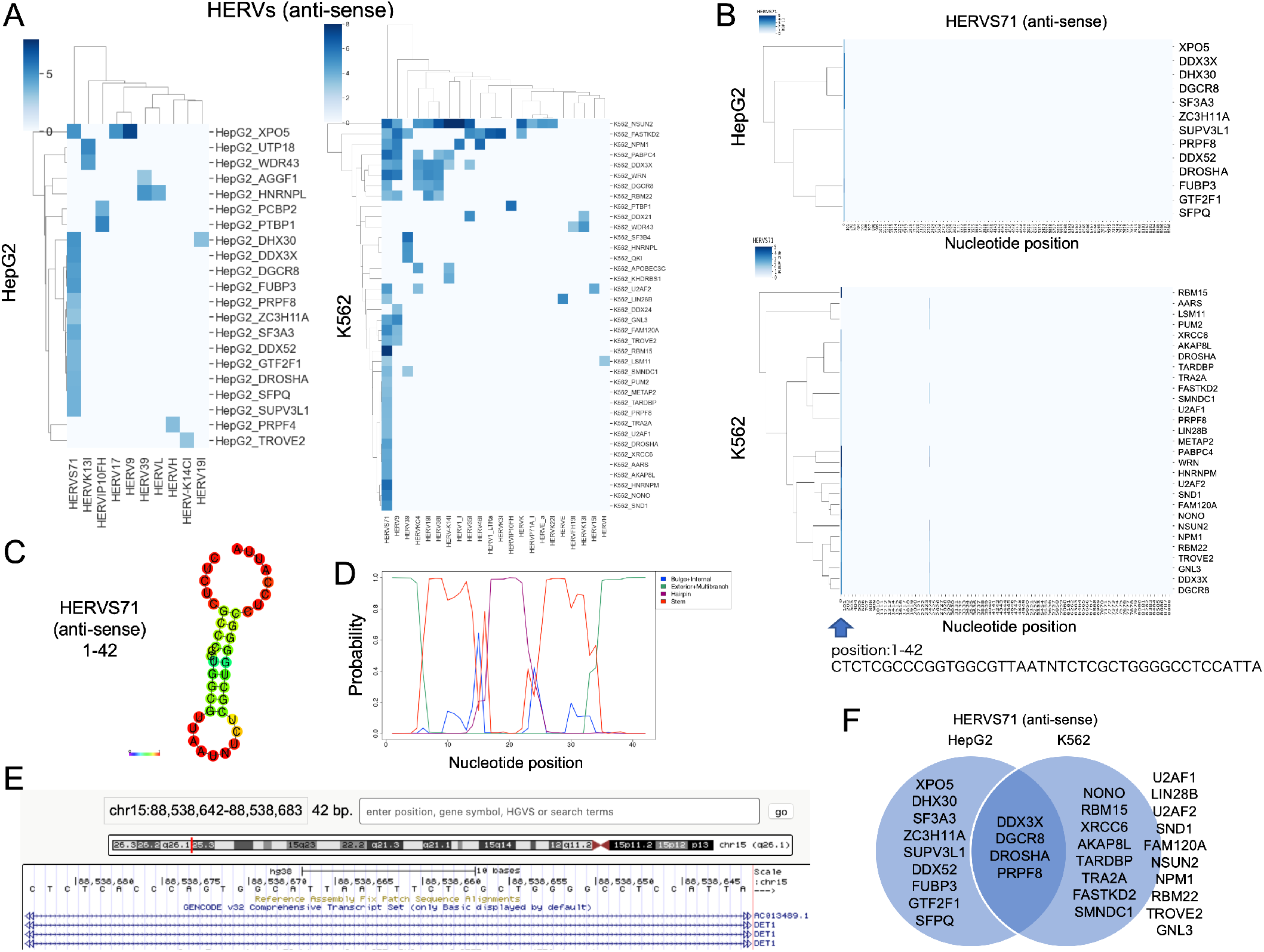
Analysis of RBP binding patterns on anti-sense of HERV subfamilies with nucleotides resolution predicts putative functional RNA elements with multiple binding RBPs. (A)RBP binding pattern of the anti-sense strand of HERV subfamilies. The x-axis indicates anti-sense of HERV subfamilies, and the y-axis indicates RBPs. Blue colors indicate the maximum values of log2 fold change of eCLIP IP signals compared to SMInput. (B)RBP binding pattern of the HERVS71 anti-sense strand with nucleotide resolution. The x-axis indicates the nucleotide position of anti-sense HERVS71 and the y-axis indicates RBPs. Blue colors indicate log2 fold change of eCLIP IP signals compared to SMInput.(C)RNA secondary structure prediction of RBP cluster sites of HERVS71(1-42) calculated by CentroidFold. Colored bases indicate base pairing probabilities. (D)Probability of RNA secondary structure of the anti-sense sequence of HERVS71(1-42) calculated by CapR. The x-axis indicates the nucleotide position of anti-sense HERVS71(1-42) and the y-axis indicates the probability of the structural profiles. (E)Example of anti-sense HERVS71(1-42) in intron of gene. HERVS71(1-42) sequence in the first intron of DET1 gene is indicated in the UCSC genome browser (hg38). (F)Venn diagram for RBP binding to the anti-sense of the HERVS71 subfamily.

## DISCUSSION

### Repeat element as an archetype of RNA funtional element

We re-analyzed RBP binding patterns on human repeat sequences, including transposons, using ENCODE eCLIP data. This analysis revealed a novel pattern of RBP binding on the human repeat sequences and putative RBP complexes that bind to repeat-derived RNAs. Notably, we found putative novel components of RBP complexes that regulate the expression of LINE1 sense strand (see Figure2). We also found that the 3’UTR of L1PA subfamilies contains putative functional elements that could be related to heterochromatin formation (see Figure4). Moreover, we found several new candidates that are components of the splicing complex that bind to the LINE1 anti-sense sequence, as well as a candidate sequence for the RBP binding in the same sequence (see Figure5). Therefore, this approach promises to identify putative functional RNA elements, which are derived from repeat sequences including transposons. Previous reports have not focused on the precise binding patterns of RBPs to the repeat sequences, especially to TE and its subfamily (24). In contrast, we focused on RBP binding patterns to TEs with nucleotide resolution and found several short RNA fragments that bind to multiple RBPs and form RBP clusters (see Figure2-7). Taking advantage of eCLIP data, which detects precise binding sites between RNA and protein, we predicted RBP binding sites and RNA secondary structures of the short RNA fragments and found that these sequences can form stable stem-loop structures. Because TE sequences are derived from ancient viral sequences, they often contain many repetitive sequences that form specific RNA secondary structures. These elements are thought to act as key functional elements for the TE regulation (45). We found that such fragments can bind to multiple endogenous RBPs and form an RBP cluster, suggesting that these elements may also have some function in the host transcriptome regulation. Indeed, previous analysis of RBP binding to TEs concluded that TE fragments are an important target of RBPs (64). Thus, it is quite possible that these short RNA fragments survived natural selection and were co-opted by the host genome during evolution. We propose that TE sequences contain many short fragments, which are the archetypes of RNA functional elements for both TEs and the host genome. Consistent with this idea, a recent report has revealed that one of the functional elements of Xist, A-repeat, is derived from a TE sequence (9). This case is considered to be the result of adaptation between the TE and the host genome. Similar cases were considered in other TEs and were proposed as an RIDL hypothesis (65). In addition to the cases of lncRNAs, this concept can be applied to the splicing regulation of intronic TE elements. Our analysis and data from other studies suggest that the L1 sequence is a very important element for splicing (5, 24, 64, 66). Thus, our study will contribute to finding new functional elements in other biological contexts. The next question is whether the ncRNAs, which contain repeat-derived putative functional elements provided as candidates in this study, actually bind to predicted RBPs because our analysis only predicts the binding of repeat RNAs to specific RBPs. In addition, it is necessary to confirm whether these sequences contribute to the function of the corresponding repeat-containing RNAs such as lncRNA, introns, or UTRs.

### Co-evolution of RNA functional elements and TEs

Interestingly, TE sequences showed different and specific RBP binding patterns among subfamilies and strands, suggesting that each of the fragments might have distinct functions. We cannot identify whether all of these sequences have associated functions. However, it is possible that each of these sequences acts as functional RNA elements in some contexts, since our analysis successfully confirmed the functional complex on sense and anti-sense strands of LINE1 sequences (see Figure2-5). In addition, we found some examples of repeat-derived fragments that bind to multiple RBPs in HERV subfamilies (see Figure6, 7). Importantly, different subfamilies of LINE1 seem to have different RBP binding patterns with different functions (5), suggesting that functional RNA elements might have evolved along with TEs and were selected during evolution. Indeed, our study revealed that L1PA4-8, a group of L1PA subfamilies, had distinct RBP binding patterns that included SAFB, SAFB2, and HNRNPL from other subfamilies in a short fragment within the 3’UTR (see Figure 4). Because these RBPs have a repressive function for transcription, L1PA4-8 might be inhibited by this RBP complex, whereas younger subfamilies such as L1HS might have undergone some mutations to escape this complex. Thus, fragments derived from L1PA4-8 might become a scaffold for the transcriptionally repressive RBP complex in introns of host genes. This scenario could explain how the TE-derived sequence became a new functional element. Thus, we hypothesize that an arms race between LINE1 and host immunity could be a source of creating RNA functional elements and promoting co-evolution.

### Repeat-derived sequence and liquid-liquid phase separation (LLPS)

Given that merged eCLIP data represent not necessarily simultaneous binding of RBPs at specific target RNA, we cannot determine which proteins are binding the same position at the same time. However, we consider the possibility that particular RBP complexes bind to specific RNA sequences simultaneously through mechanisms such as LLPS, which enables the association of specific RBPs and RNA sequences with each other in the same droplet (67). A recent report has shown that the SAFB complex with repeat-derived RNAs, including LINEs, can form LLPS droplets and contribute to heterochromatin formation (50). Furthermore, other groups have also shown that LINE1 RNA sequences are associated with transcriptionally inactive domains and are essential for chromatin architecture (68, 69, 70, 71). However, which part of the LINE1 sequence is crucial for LLPS formation and subsequent function remains elusive. In this study, we identified several candidates for the functional elements of LINE subfamilies that harbor multiple binding sites for RBPs. We also predicted that these fragments could form stable secondary RNA structures. Notably, both the stoichiometry of RBP binding sites and RNA secondary structures could be essential for LLPS (72, 73). Therefore, our data provide promising candidates for functional RNA elements derived from TE sequences. We believe our predictions of functional elements in repeat RNAs using eCLIP data provide new insights into RBP binding mechanisms, including LLPS, to repeat sequences.

## CONCLUSION

We focused on the repeat sequences on RNA and found that many RBPs bind to these elements. Our analysis revealed that at least some of these elements form stable RNA secondary structures and are associate with multiple RBPs. Overall, we found several RNA fragments that are the strong candidates for RNA functional elements. Further investigation is required to determine the functions associated with these sequences.

## ACKNOWLEDGEMENTS

We thank Mr. Yu Hamaguchi for technical information and assistance, Dr. Martin Frith, Dr. Tsukasa Fukunaga, and Dr. Yutaka Saito for fruitful discussions. Computation for this study was partially performed on the NIG supercomputer at ROIS National Institute of Genetics.

