## Supplemental Figures for "Binding patterns of RNA binding proteins to repeat-derived RNA sequences reveal putative functional RNA elements"

Supplementary Figure 1

A

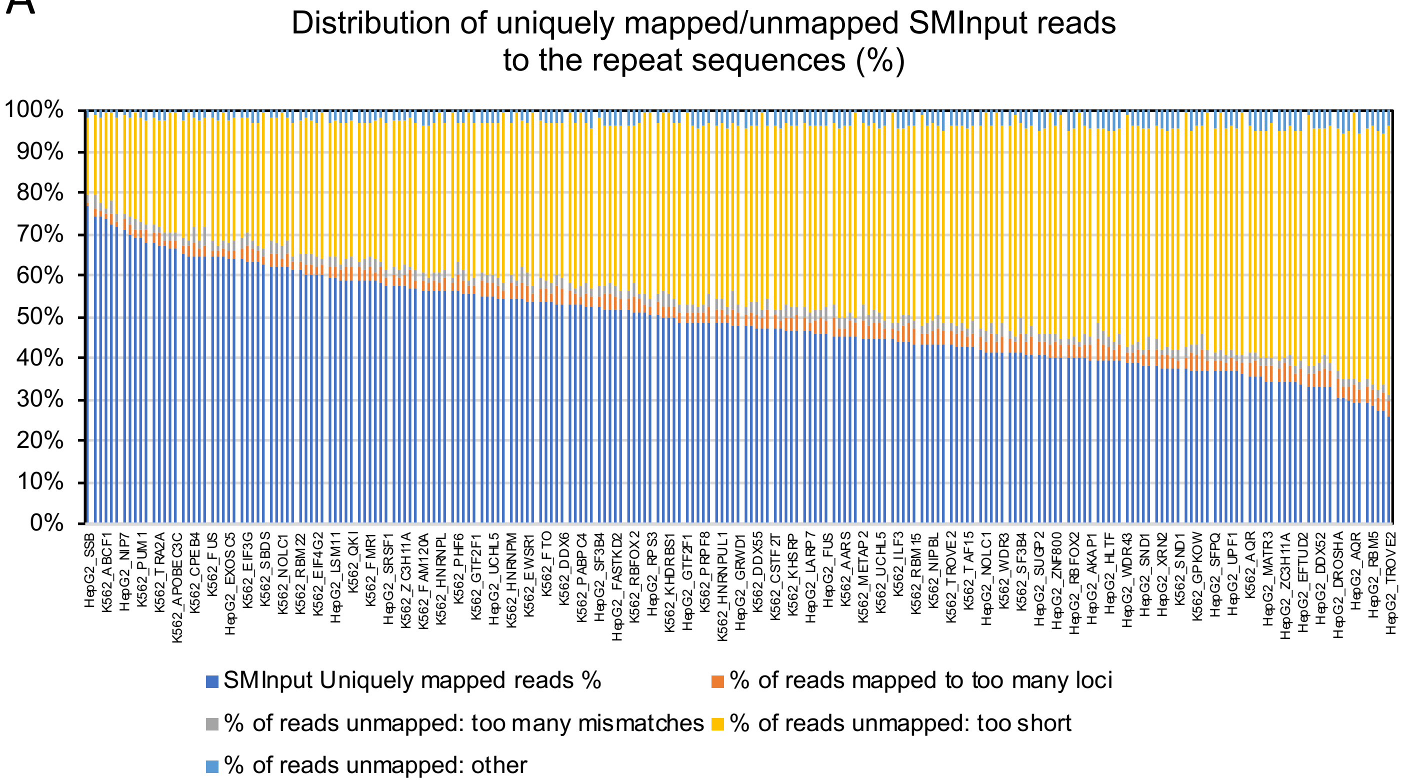


B

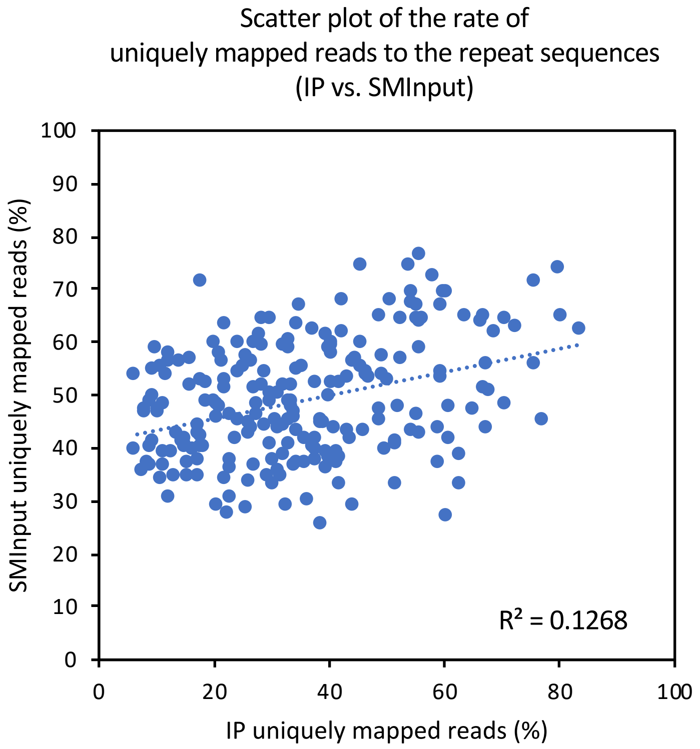


Supplementary Figure 1

(A) Distribution of uniquely mapped or unmapped SMInput reads of each RBP to the repeat sequences in HepG2 or K562 cells. The x-axis indicates RBPs, and the y-axis indicates the percentage of mapped/unmapped reads.

(B) Scatter plot of the rate of uniquely mapped reads to the repeat sequences (IP vs. SMInput). The x-axis indicates the percentage of uniquely mapped IP reads, and the y-axis indicates the percentage of uniquely mapped SMInput reads. Blue dots represent the RBPs.

Supplementary Figure 1

C

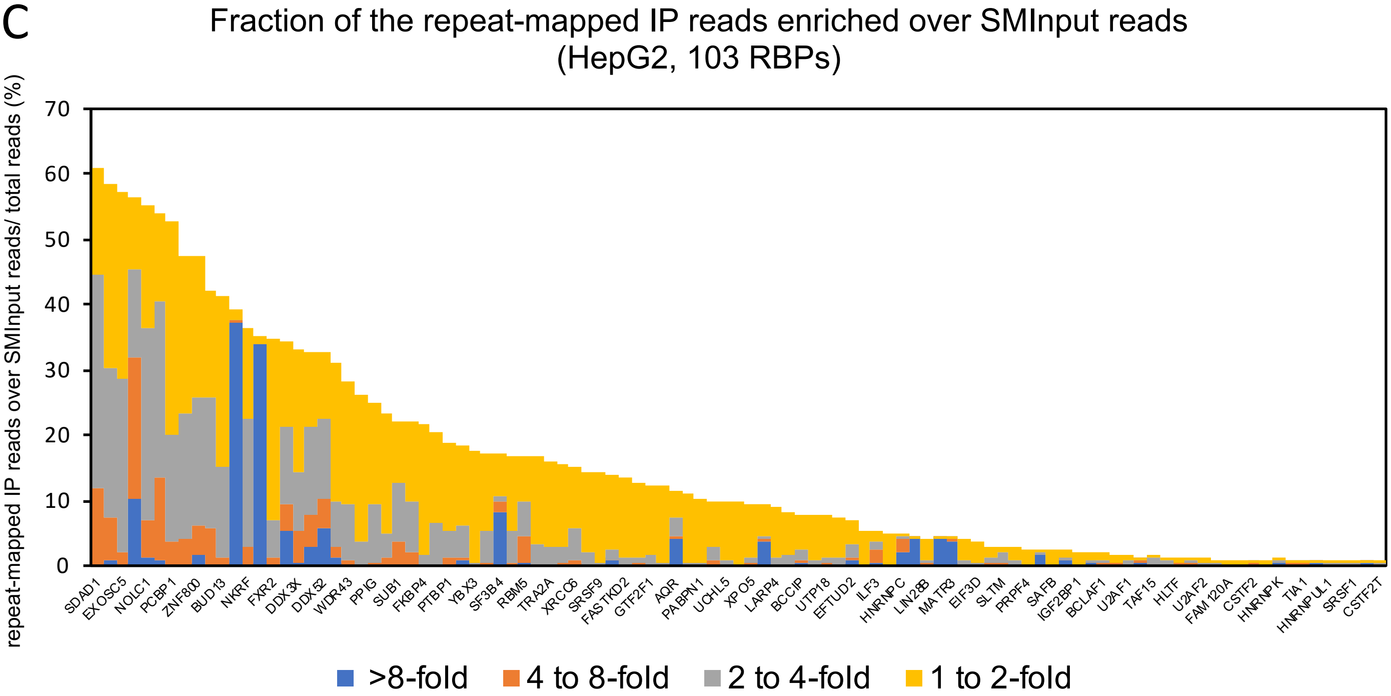


D

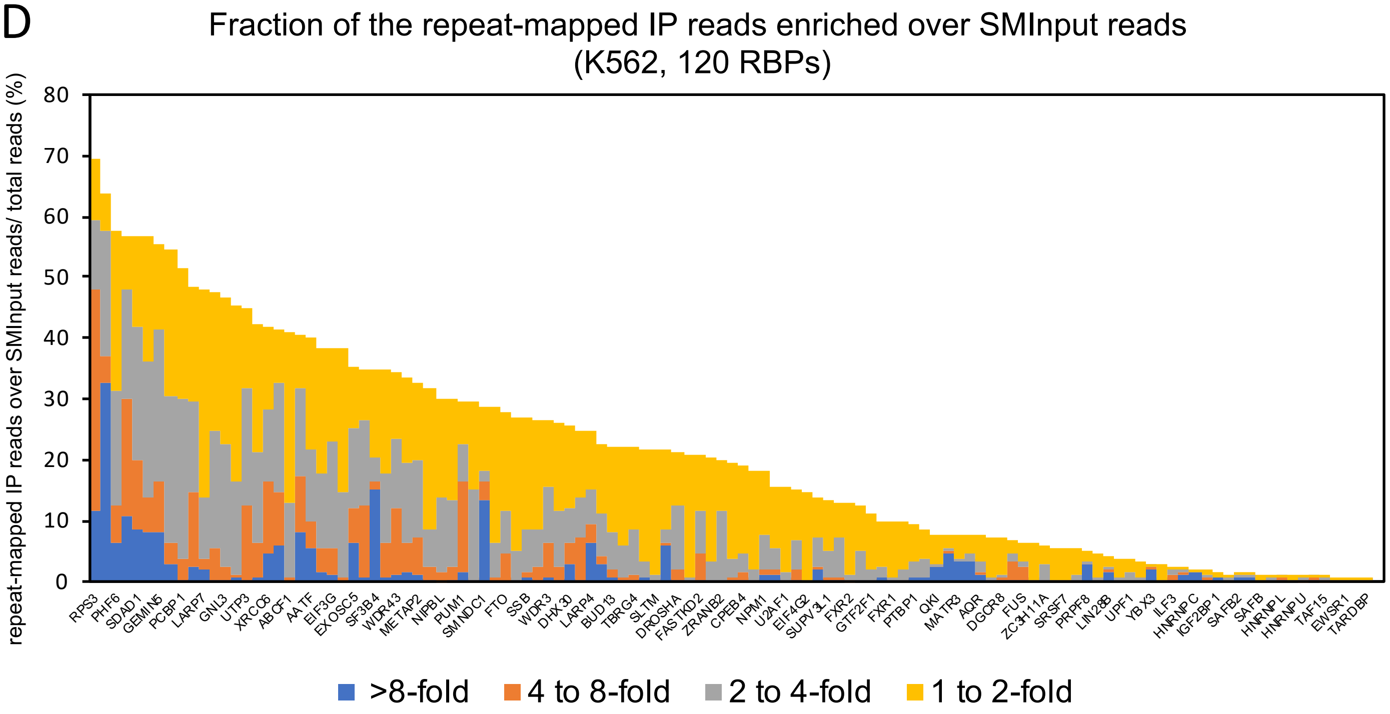


E

| | >8-fold | >4-fold | >2-fold | >1-fold |
|---|---|---|---|---|
| K562 average (%) | 1.86 | 4.47 | 10.75 | 20.95 |
| HepG2 average (%) | 1.37 | 2.69 | 7.04 | 15.35 |

Supplementary Figure 1
(C,D) Fraction of repeat-mapped IP reads enriched over SMInput reads of HepG2 (C) or K562 (D) cells. The x-axis indicates RBPs, and the y-axis indicates the percentage of repeat-mapped IP reads enriched over SMInput in the total IP reads. Each of the enrichment rates (IP/SMInput) is indicated by colored bars.
(E) Average rates of IP reads enriched 1-, 2-, 4-, and 8-fold over SMInput in HepG2 and K562 cells.

Supplementary Figure 1

F

| | HepG2 top30 | K562 top30 | HepG2 bottom30 | K562 bottom30 |
|---|---|---|---|---|
| 1 | SDAD1 | RPS3 | AGGF1 | SRSF7 |
| 2 | NOL12 | RPS11 | PRPF4 | PUM2 |
| 3 | EXOSC5 | PHF6 | QKI | PRPF8 |
| 4 | RPS3 | DDX24 | SAFB | RBFOX2 |
| 5 | NOLC1 | SDAD1 | LSM11 | LIN28B |
| 6 | DKC1 | DDX51 | IGF2BP1 | NONO |
| 7 | PCBP1 | GEMIN5 | DGCR8 | UPF1 |
| 8 | FTO | SBDS | BCLAF1 | SRSF1 |
| 9 | ZNF800 | PCBP1 | GRWD1 | YBX3 |
| 10 | NIP7 | DDX52 | U2AF1 | HNRNPUL1 |
| 11 | BUD13 | LARP7 | PCBP2 | ILF3 |
| 12 | SF3A3 | ZNF800 | TAF15 | PABPC4 |
| 13 | NKRF | GNL3 | HNRNPU | HNRNPC |
| 14 | SMNDC1 | DDX42 | HLTF | TIA1 |
| 15 | FXR2 | UTP3 | HNRNPA1 | IGF2BP1 |
| 16 | LARP7 | DDX21 | U2AF2 | HNRNPA1 |
| 17 | DDX3X | XRCC6 | RBFOX2 | SAFB2 |
| 18 | TROVE2 | APOBEC3C | FAM120A | KHDRBS1 |
| 19 | DDX52 | ABCF1 | TIAL1 | SAFB |
| 20 | G3BP1 | ZC3H8 | CSTF2 | FAM120A |
| 21 | WDR43 | AATF | ZC3H11A | HNRNPL |
| 22 | AKAP1 | FMR1 | HNRNPK | U2AF2 |
| 23 | PPIG | EIF3G | FUBP3 | HNRNPU |
| 24 | DDX55 | MTPAP | TIA1 | HNRNPK |
| 25 | SUB1 | EXOSC5 | SRSF7 | TAF15 |
| 26 | EIF3H | NCBP2 | HNRNPUL1 | KHSRP |
| 27 | FKBP4 | SF3B4 | SFPQ | EWSR1 |
| 28 | DDX6 | PUS1 | SRSF1 | CSTF2T |
| 29 | PTBP1 | WDR43 | HNRNPL | TARDBP |
| 30 | POLR2G | WRN | CSTF2T | SERBP1 |

G

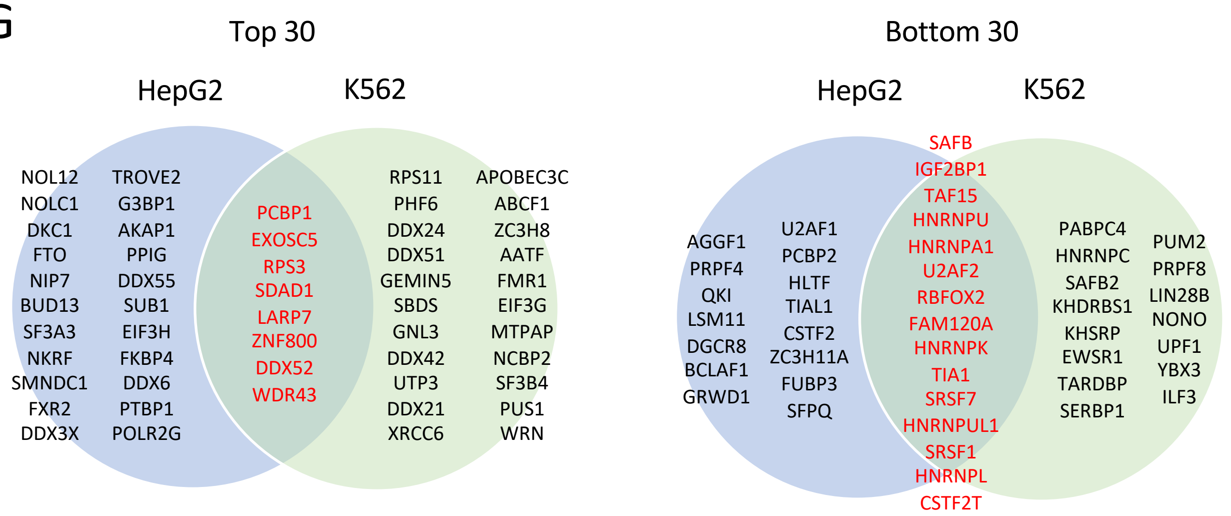


Supplementary Figure 1

(F) List of the top or bottom 30 repeat-associated RBPs in HepG2 or K562 cells. The fraction of repeat-associated IP reads that contribute to RBP binding events for each RBP were calculated in (C, D). The common RBPs between HepG2 and K562 cells are indicated by red letters.

(G) Venn diagram of the top or bottom 30 repeat associated RBPs in HepG2 and K562 cells.

Supplementary Figure 1

H

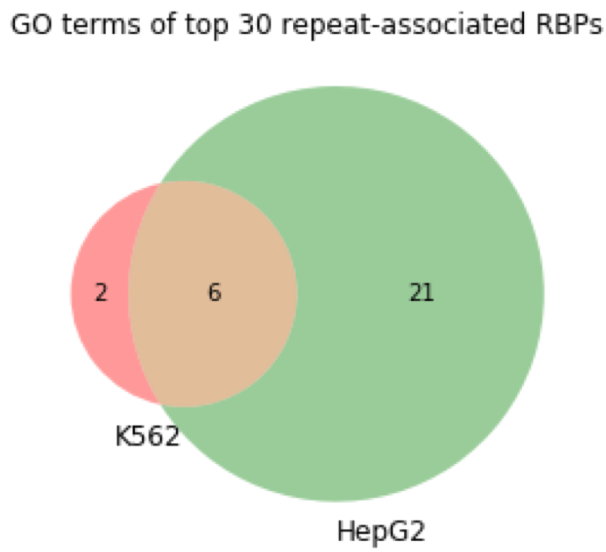


I

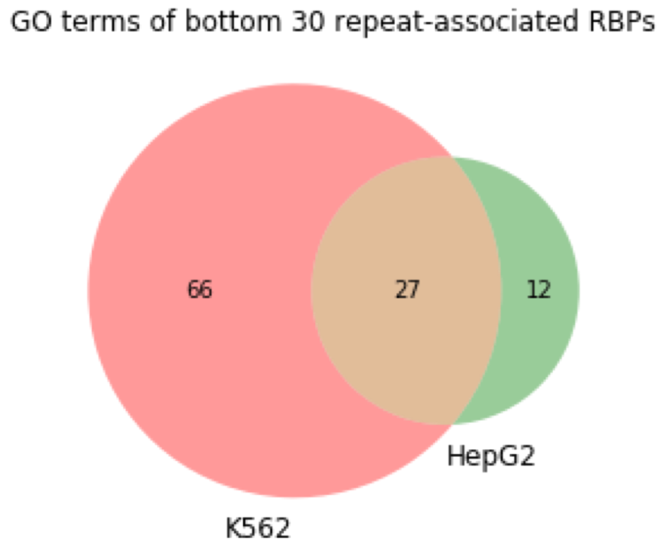


Supplementary Figure 1
(H, I) Venn diagram for GO terms of the top (H) or bottom (I) 30 of repeat-associated RBPs in K562 and HepG2 cells. The numbers in circles indicate numbers of specifically enriched GO terms in the top (H) or bottom (I) 30 of repeat-associated RBPs in K562 or HepG2 cells.

Supplementary Figure 1

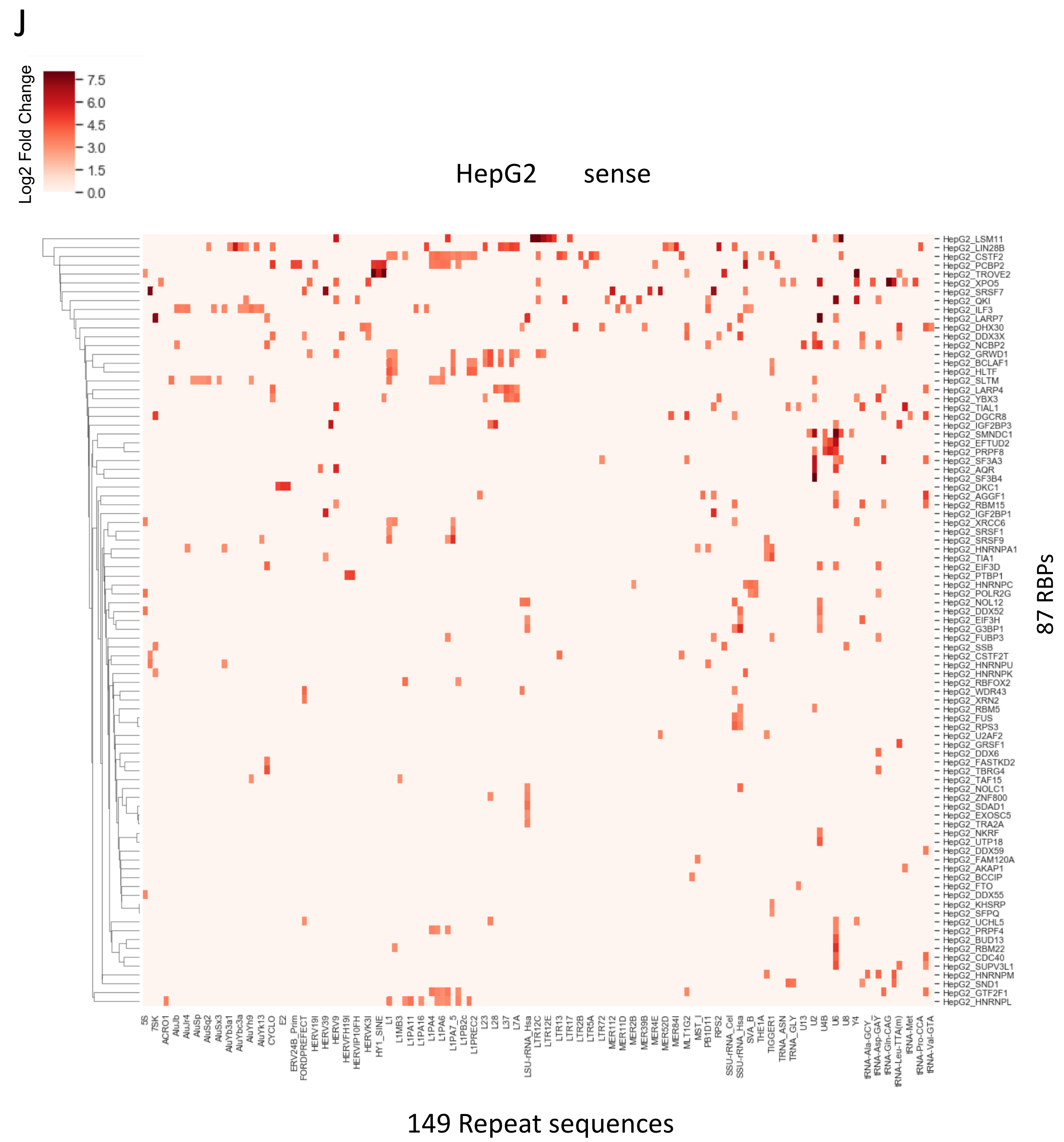


Supplementary Figure 1
(J) RBP binding pattern of the sense strand of repeat sequences in HepG2 cells. The x-axis shows repeat sequences and the y-axis shows RBPs. Red colors in the heat-map indicate the maximum values of log2 fold change of eCLIP IP signals compared to SMInput.

Supplementary Figure 1

K

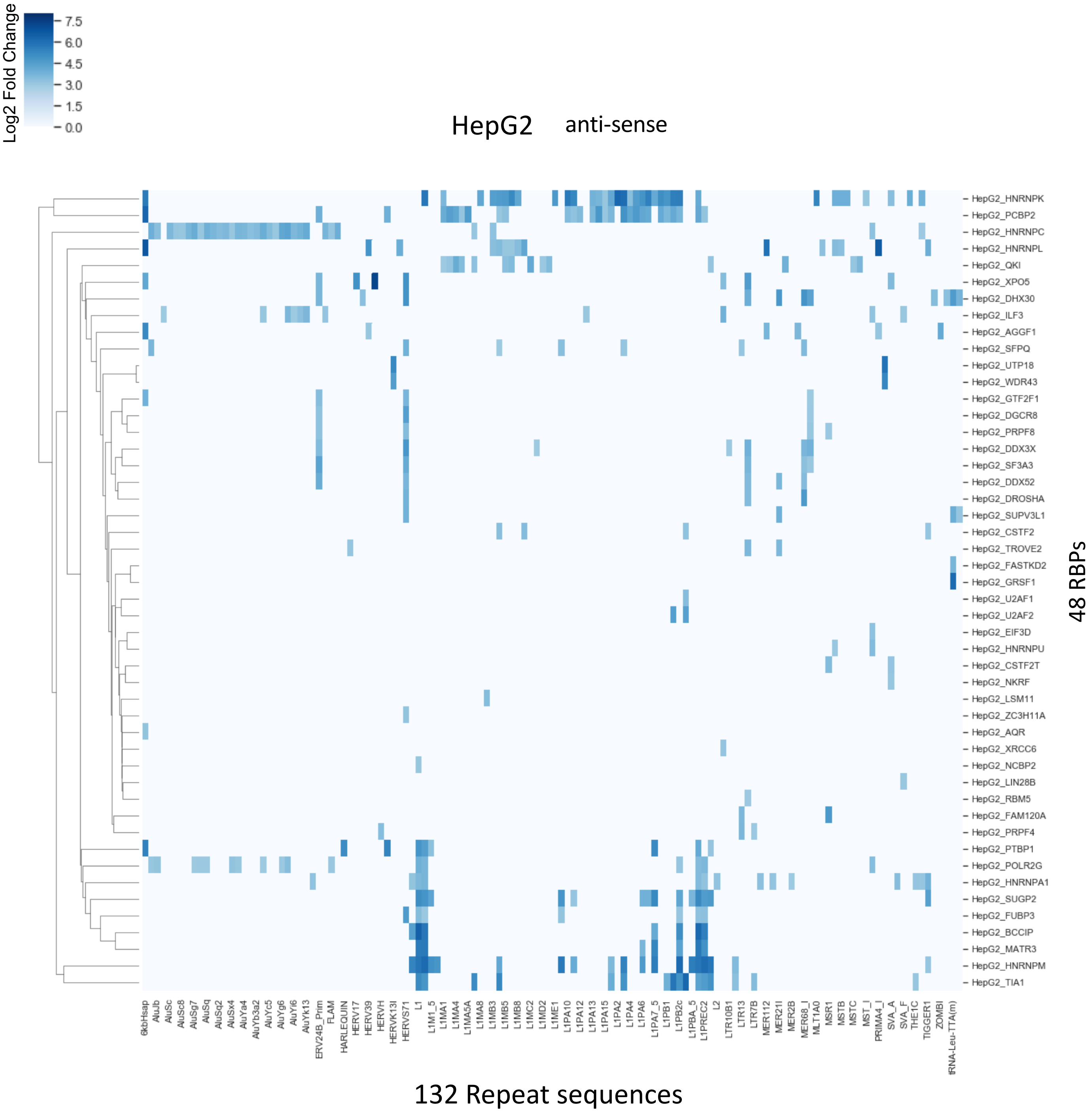


Supplementary Figure 1

(K) RBP binding pattern of the anti-sense strand of repeat sequences in HepG2 cells. The x-axis shows repeat sequences and the y-axis shows RBPs. Blue colors in the heat-map indicate the maximum values of log2 fold change of eCLIP IP signals compared to SMInput.

Supplementary Figure 1

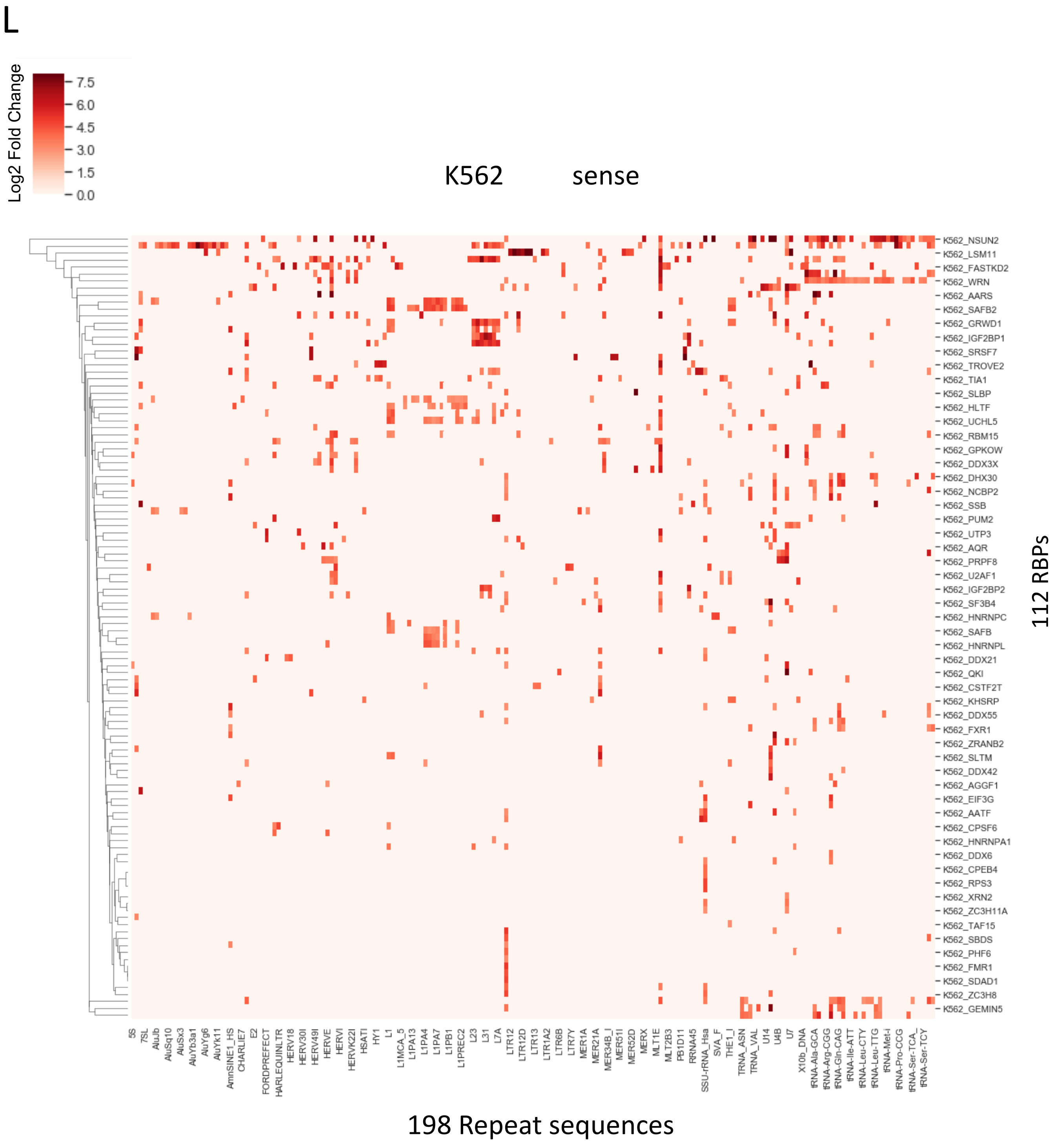


Supplementary Figure 1
(L) RBP binding pattern of the sense strand of repeat sequences in K562 cells. The x-axis shows repeat sequences and the y-axis shows RBPs. Red colors in the heat-map indicate the maximum values of log2 fold change of eCLIP IP signals compared to SMInput.

#### Supplementary Figure 1

M

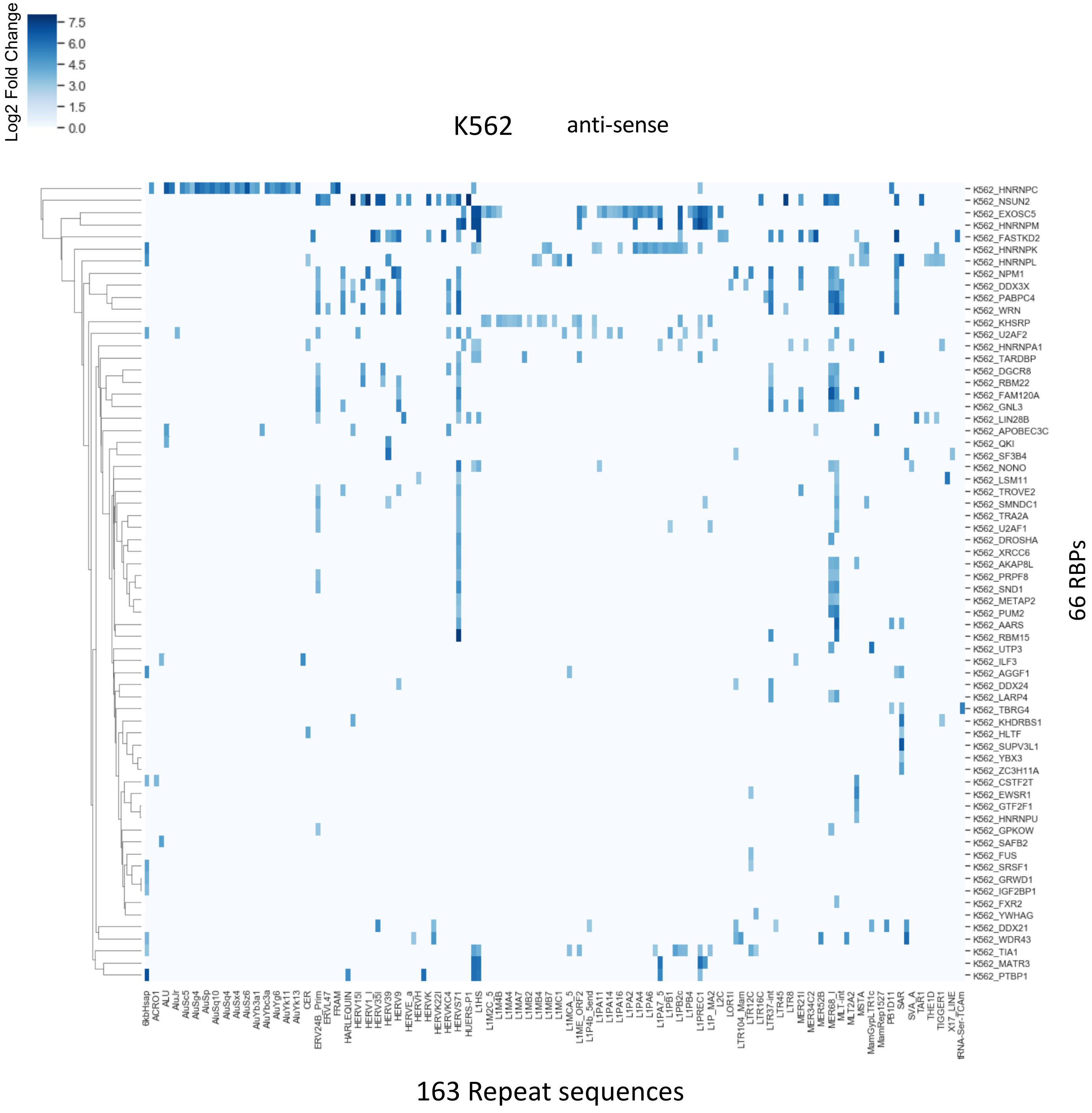


Supplementary Figure 1
(M) RBP binding pattern of the anti-sense strand of repeat sequences in K562 cells. The x-axis shows repeat sequences and the y-axis shows RBPs. Blue colors in the heat-map indicate the maximum values of log2 fold change of eCLIP IP signals compared to SMInput.

### Supplementary Figure 2

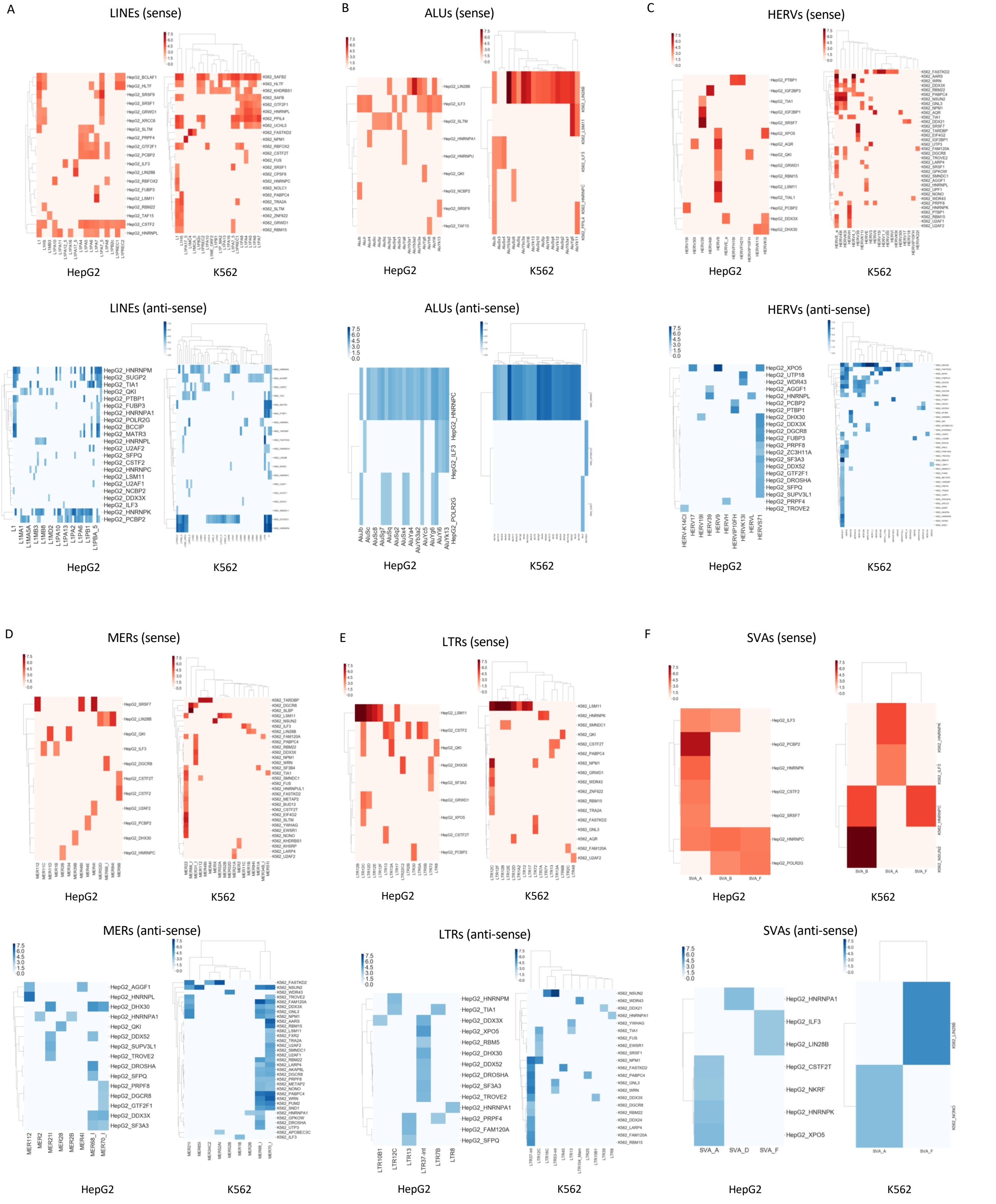


Supplementary Figure 2
RBP binding patterns of TE subfamilies (A:LINE, B:ALU, C:HERV, D:MER, E:LTR, F:SVA). The x-axis indicate sense or anti-sense strand of TE subfamilies and the y-axis indicate RBPs. Red (sense strand) or blue (anti-sense strand) colors indicate the maximum values of log2 fold change of eCLIP IP signals compared to SMInput.

Supplementary Figure 3

A

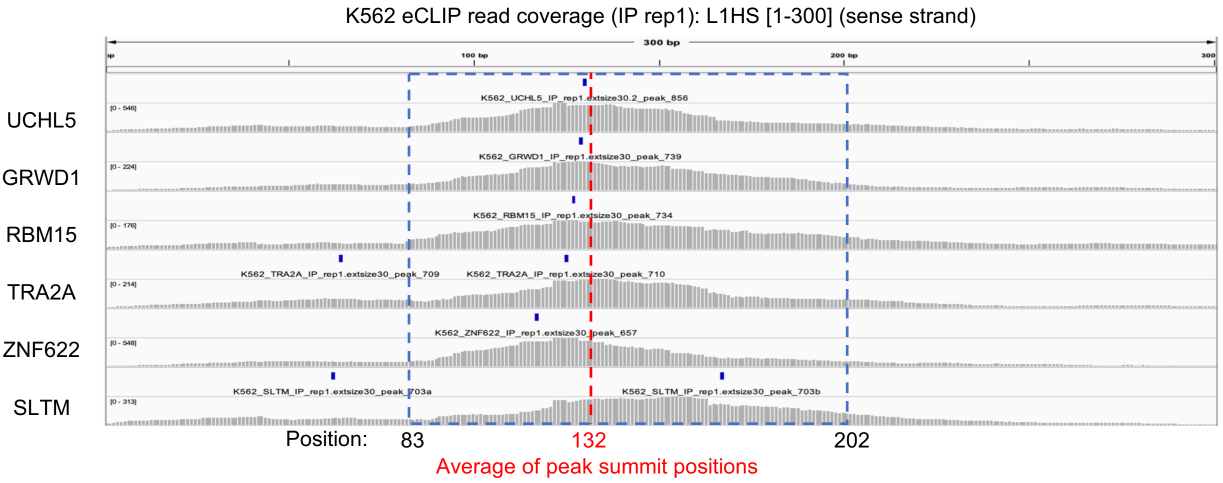


B

>L1HS [83-202] (120 nt)
CTGCATTTCCATCTGAGGTACCGGGTTCATCTCACTAGGGAGTGCCAGACAGTGGGCGCAGGCCAGTGTGTGTGCGCACCGTGCGCGAGCCGAAGCAGGGCGAGGCATTGCCTCACCTGG

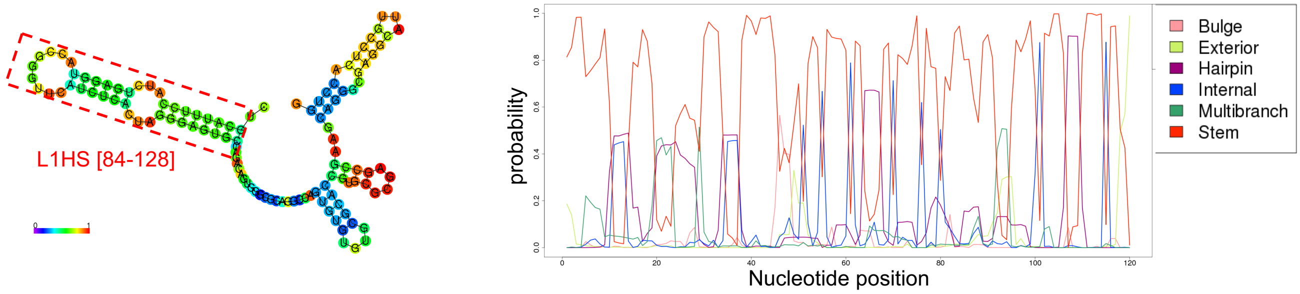


C Upstream of peak summit average [~132]

> L1HS [93-132] (40 nt)
ATCTGAGGTACCGGGTTCATCTCACTAGGGAGTGCCAGAC

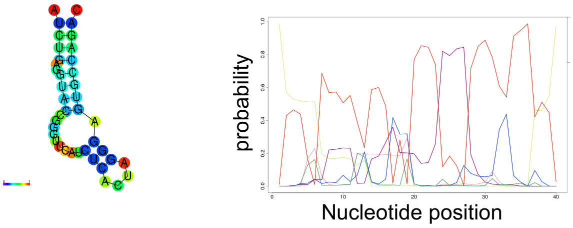


>L1HS [83-132] (50 nt)
CTGCATTTCCATCTGAGGTACCGGGTTCATCTCACTAGGGAGTGCCAGAC

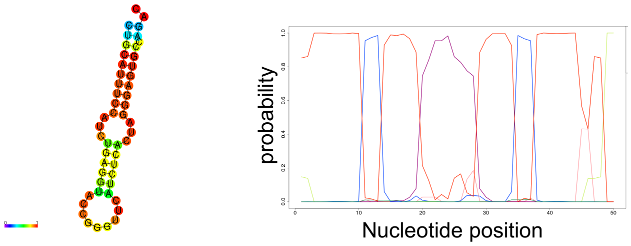


> L1HS [63-132] (70 nt)
ACGCAGAAGACGGGTGATTTCTGCATTTCCATCTGAGGTACCGGGTTCATCTCA
CTAGGGAGTGCCAGAC

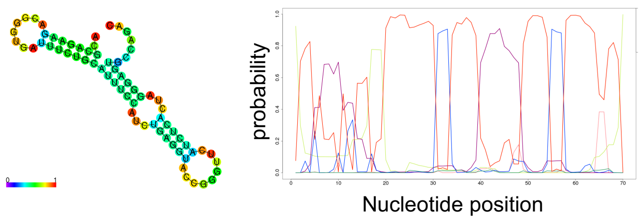


D Downstream of peak summit average [133~]

>L1HS [133-172] (40 nt)
AGTGGGCGCAGGCCAGTGTGTGTGCGCACCGTGCGCGAGC

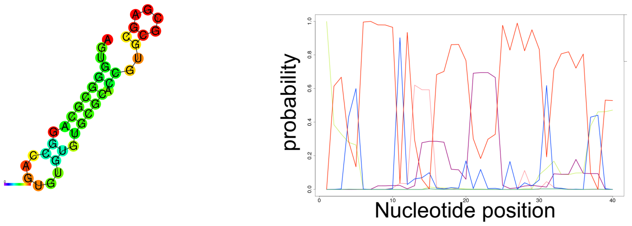


>L1HS [133-182] (50 nt)
AGTGGGCGCAGGCCAGTGTGTGTGCGCACCGTGCGCGAGCCGAAGCAGGG

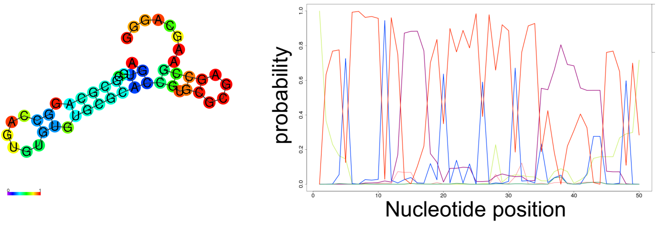


>L1HS [133-202] (70 nt)
AGTGGGCGCAGGCCAGTGTGTGTGCGCACCGTGCGCGAGCCGAAGCAGGGCG
AGGCATTGCCTCACCTGG

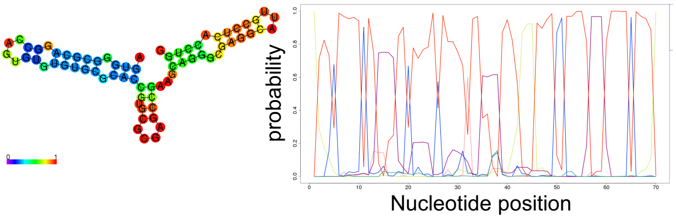

Supplementary Figure 3

E

K562 eCLIP read coverage (IP rep1): L1HS [4480-4620] (sense strand)

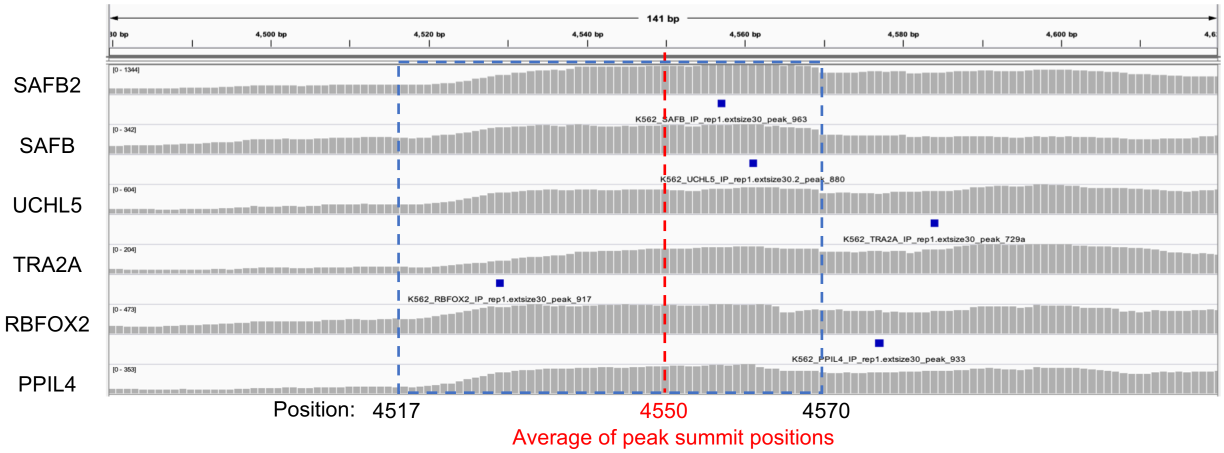


F

>L1HS [4517-4570] (54 nt)
ACTTTAAAGTTCATATGGAACCAAAAAAGAGCCCGCATCGCCAAGTCAATCCTA

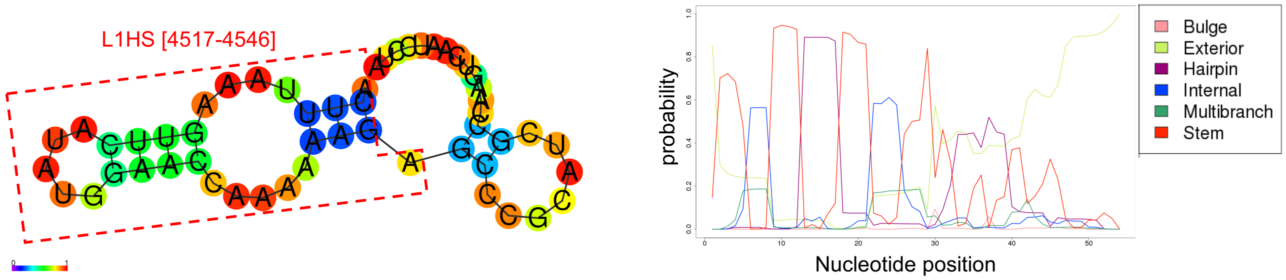


G Upstream of peak summit average [~4550]

>L1HS [4521-4550] (30 nt)
TAAAGTTCATATGGAACCAAAAAAGAGCCC

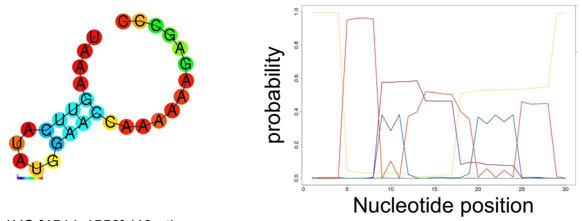


>L1HS [4511-4550] (40 nt)
AAAACTACTTTAAAGTTCATATGGAACCAAAAAAGAGCCC

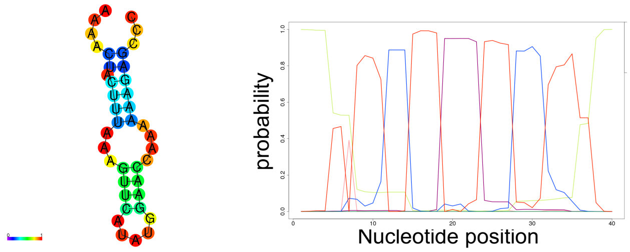


>L1HS [4501-4550] (50 nt)
AGAATTGGAAAAAACTACTTTAAAGTTCATATGGAACCAAAAAAGAGCCC

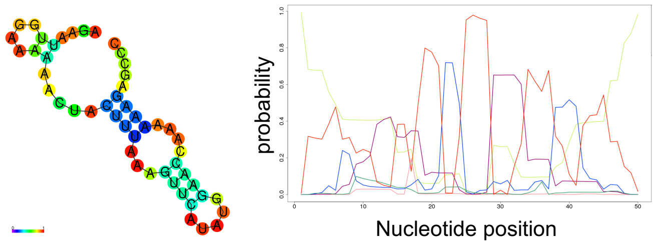


H Downstream of peak summit average [4551~]

>L1HS [4551-4580] (30 nt)
GCATCGCCAAGTCAATCCTAAGCCAAAAGA

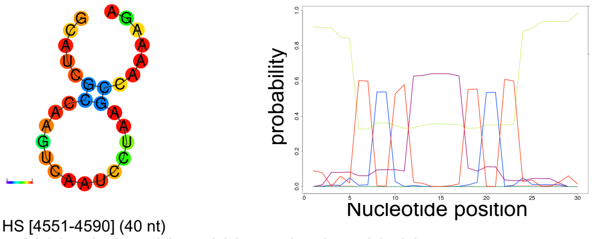


>L1HS [4551-4590] (40 nt)
GCATCGCCAAGTCAATCCTAAGCCAAAAGAACAAAGCTGG

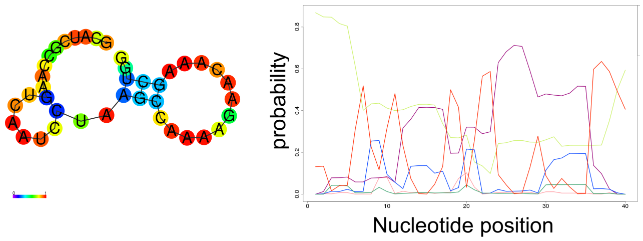


>L1HS [4551-4600] (50 nt)
GCATCGCCAAGTCAATCCTAAGCCAAAAGAACAAAGCTGGAGGCATCACA

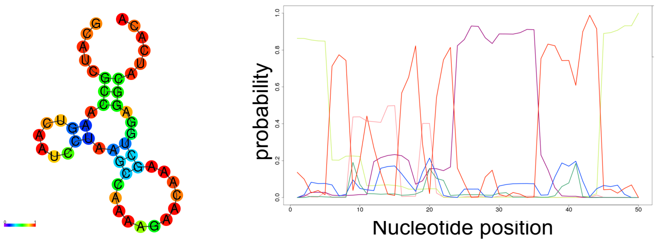

Supplementary Figure 3
(A) eCLIP IP read coverage of RBPs associated with L1HS [1-300] in K562 cells. The y-axis shows RBP names, and the x-axis shows the nucleotide position of the L1HS sequence. The red vertical bar indicates the average of the peak summit positions of RBPs indicated in the IGV tracks. The blue rectangle indicates the RBP binding sites of L1HS[83-202] covered with eCLIP IP reads. (B,C,D) Prediction of RNA secondary structures for RBP cluster sites for 40 nt, 50 nt, and 70 nt upstream (C) or 40 nt, 50 nt, and 70 nt downstream (D) or both upstream and downstream (B) regions of the peak summit average [132] calculated using CentroidFold (left panels) or CapR (right panels). Colored bases of the left panels (CentroidFold) indicate the base-pairing and loop probabilities. The right panels (CapR) show the structural profile of an RNA base for a set of six probabilities (stem part, hairpin loop, bulge loop, internal loop, multibranch loop, and exterior loop) that the base belongs to each category. The x-axis indicates the nucleotide position, and the y-axis indicates the probability of the structural profiles. The red rectangle in (B) indicates L1HS[84-128] that was predicted to form a hairpin loop structure (see Figure3B).

Supplementary Figure 3
(E) eCLIP IP read coverage of RBPs that are associated with L1HS [4880-4620] in K562 cells. The y-axis shows the names of RBPs, and the x-axis shows the nucleotide position of the L1HS sequence. The red vertical bar indicates the average of the peak summit positions of RBPs indicated by the IGV tracks. The blue rectangle indicates the RBP binding sites of L1HS[4517-4570] covered with eCLIP IP reads. (F,G,H) Prediction of RNA secondary structures for RBP cluster sites for 30 nt, 40 nt, and 50 nt upstream(C) or 30 nt, 40 nt, and 50 nt downstream(D) or both upstream and downstream(B) regions of the peak summit average [4550] calculated using CentroidFold (left panels) or CapR (right panels). Colored bases of the left panels (CentroidFold) indicate the base-pairing and loop probabilities. The right panels (CapR) show the structural profile of an RNA base for a set of six probabilities (stem part, hairpin loop, bulge loop, internal loop, multibranch loop, and exterior loop) that the base belongs to each category. The x-axis indicates the nucleotide position, and the y-axis indicates the probability of the structural profiles. The red rectangle in (F) indicates L1HS[4517-4546] that was predicted to form a hairpin loop structure (see Figure 3F).

### Supplementary Figure 4

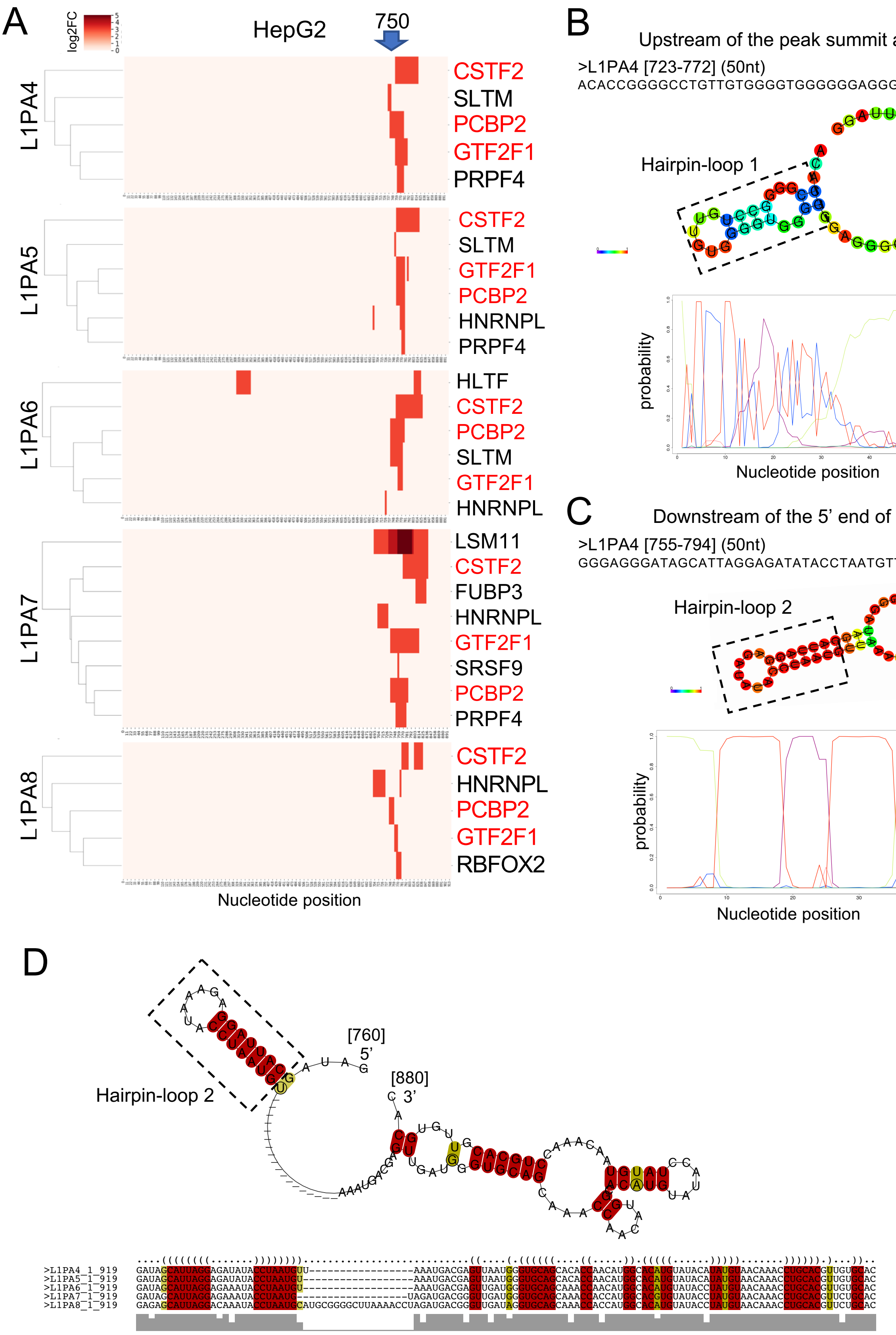


| | |
|---|---|
| Location | 760 – 880 |
| Length | 120 |
| Sequences | 5 |
| Columns | 120 |
| Reading direction | forward |
| Mean pairwise identity | 86.52 |
| Mean single sequence MFE | -27.64 |
| Consensus MFE | -21.34 |
| Energy contribution | -21.10 |
| Covariance contribution | -0.24 |
| Combinations/Pair | 1.12 |
| Mean z-score | -1.63 |
| Structure conservation index | 0.77 |
| SVM decision value | 0.23 |
| SVM RNA-class probability | 0.643315 |
| Prediction | RNA |

Supplementary Figure 4

# E

## >L1PA4/5(723-786)(64nt)

ACACCGGGGCCTGTTGTGGGGTGGGGGGAGGGGGGAGGGATAGCATTAGGAGATATACCTAATG

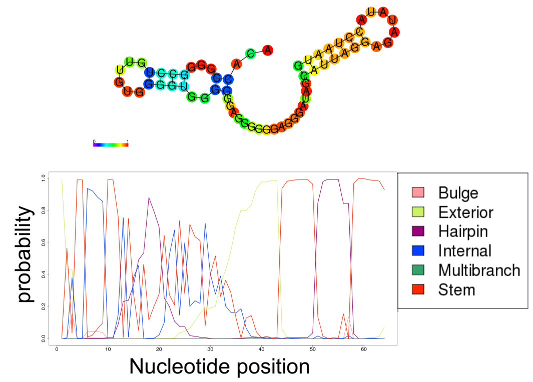


## >L1PA6(723-786)(64nt)

ACACCGGGGCCTGTCGTGGGGTGGGGGGCTGGGGGAGGGATAGCATTAGGAGAAATACCTAATG

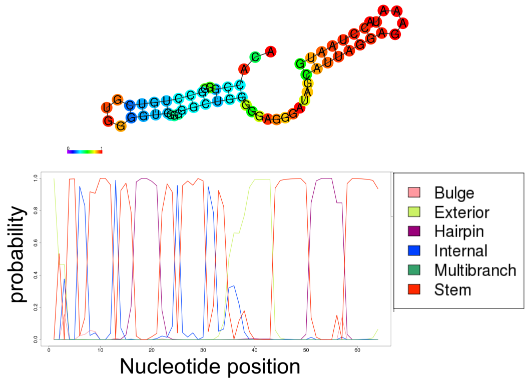


## >L1PA7(723-786)(64nt)

ACACCGGGGCCTGTCGGGGGGTGGGGGGCTAGGGGAGGGATAGCATTAGGAGAAATACCTAATG

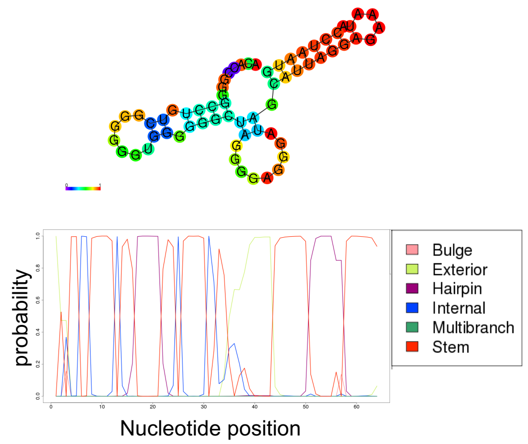


## >L1PA8(723-786)(64nt)

ACACCGGGGCCTGTCGGGGGGTGGGGGGCAAGGGGAGGGAGAGCATTAGGACAAATACCTAATG

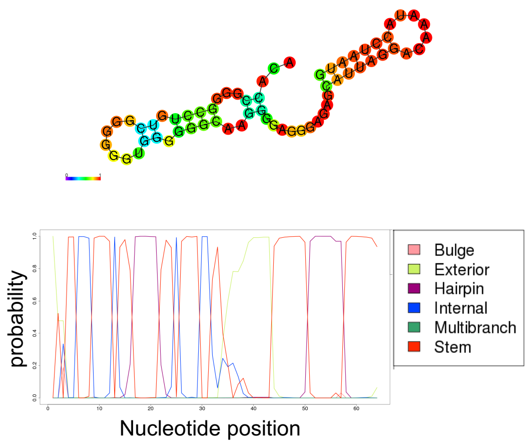


# F

## >L1PA3(723-786)(64nt)

ACTCTGGGGACTGTTGTGGGGTGGGGGGAGGGGGGAGGGATAGCATTGGGAGATATACCTAATG

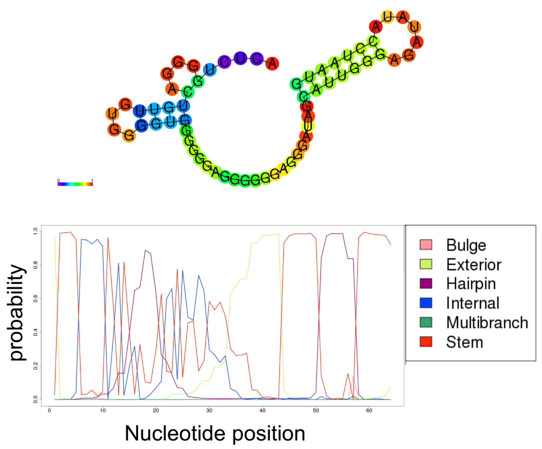


## >L1PA10(723-785)(63nt)

ACACTGGGGCCTGTCGGGGGGTGG-GGGTGGGGGGAGGGAGAGCATTAGGAAAAATAGCTAATG

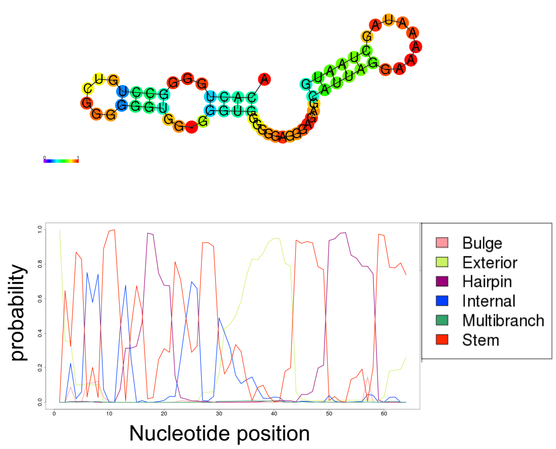

Supplementary Figure 4
(A) RBP binding pattern of L1PA4-8 sense strand (3' fragments) with nucleotide resolution in HepG2 cells. The x-axis indicates the nucleotide position of each RNA. Red colors in the heat maps indicate a log2 fold change of eCLIP IP signals compared to SMInput. RBPs written in red letters indicate common binding RBPs among L1PA4-8 in HepG2 cells. (B,C) Prediction of RNA secondary structures of L1PA4 RBP cluster sites for the upstream region of the peak summit average [723-772](B) or downstream region of the 5' end of peaks [755-794](C) calculated using CentroidFold (upper panels) or CapR (lower panels). Colored bases in the upper panels (CentroidFold) indicate the base-pairing and loop probabilities. The lower panels (CapR) show the structural profile of an RNA base for a set of six probabilities (stem part, hairpin loop, bulge loop, internal loop, multibranch loop, and exterior loop) that the base belongs to each category. The x-axis indicates the nucleotide position, and the y-axis indicates the probability of the structural profiles. (D) Prediction of consensus RNA secondary structures among L1PA4-8[760-880] using RNAz. The red and yellow letters indicate conserved base pairings with different levels of mutations. (B,C,D) The black rectangles indicate conserved hairpin loop 2 structures (see also Figure4C,D).

Supplementary Figure 4
(E,F) Prediction of RNA secondary structures of RBP cluster sites for L1PA4-8 [723-786] (E) or L1PA3 [723-786] (F, left) or L1PA10 [723-785] (F, right) calculated using CentroidFold (upper panels) or CapR (lower panels). Colored bases of the upper panels (CentroidFold) indicate the base-pairing and loop probabilities. The lower panels (CapR) show the structural profile of an RNA base for a set of six probabilities (stem part, hairpin loop, bulge loop, internal loop, multibranch loop, and exterior loop) that the base belongs to each category. The x-axis indicates the nucleotide position, and the y-axis indicates the probability of the structural profiles.

Supplementary Figure 5

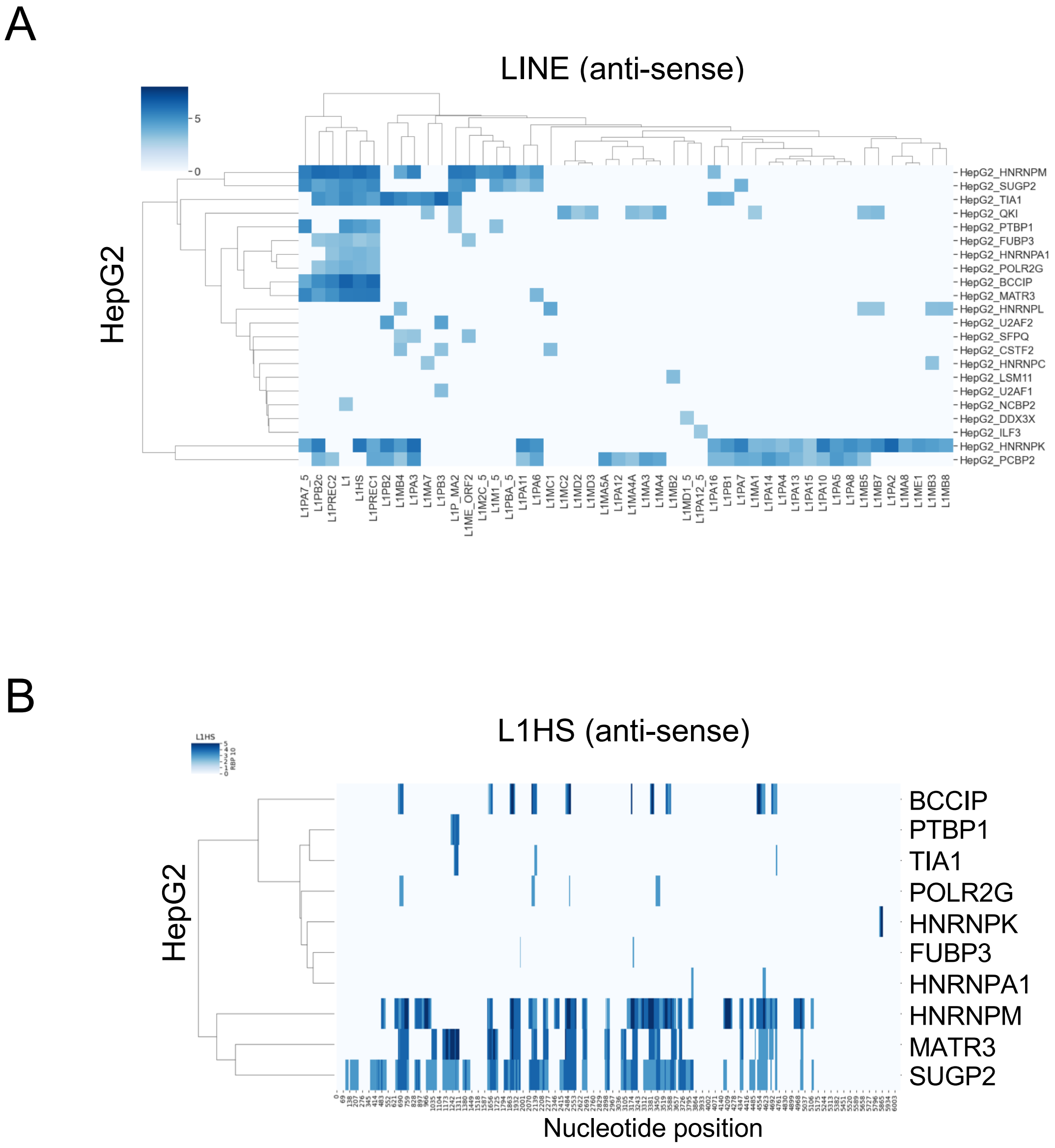


Supplementary Figure 5

(A) RBP binding pattern on anti-sense of LINE1 subfamilies in HepG2 cells. The x-axis indicates the LINE subfamilies (anti-sense strand) and the y-axis indicates RBPs. Blue colors indicate the maximum values of log2 fold change of eCLIP IP signals compared to SMInput.

(B) RBP binding pattern on anti-sense of L1HS with nucleotide resolution in HepG2 cells. The x-axis indicates the nucleotide position of the anti-sense of L1HS and y-axis indicates RBPs. Blue colors indicate log2 fold change of eCLIP IP signals compared to SMInput.

Supplementary Figure 5

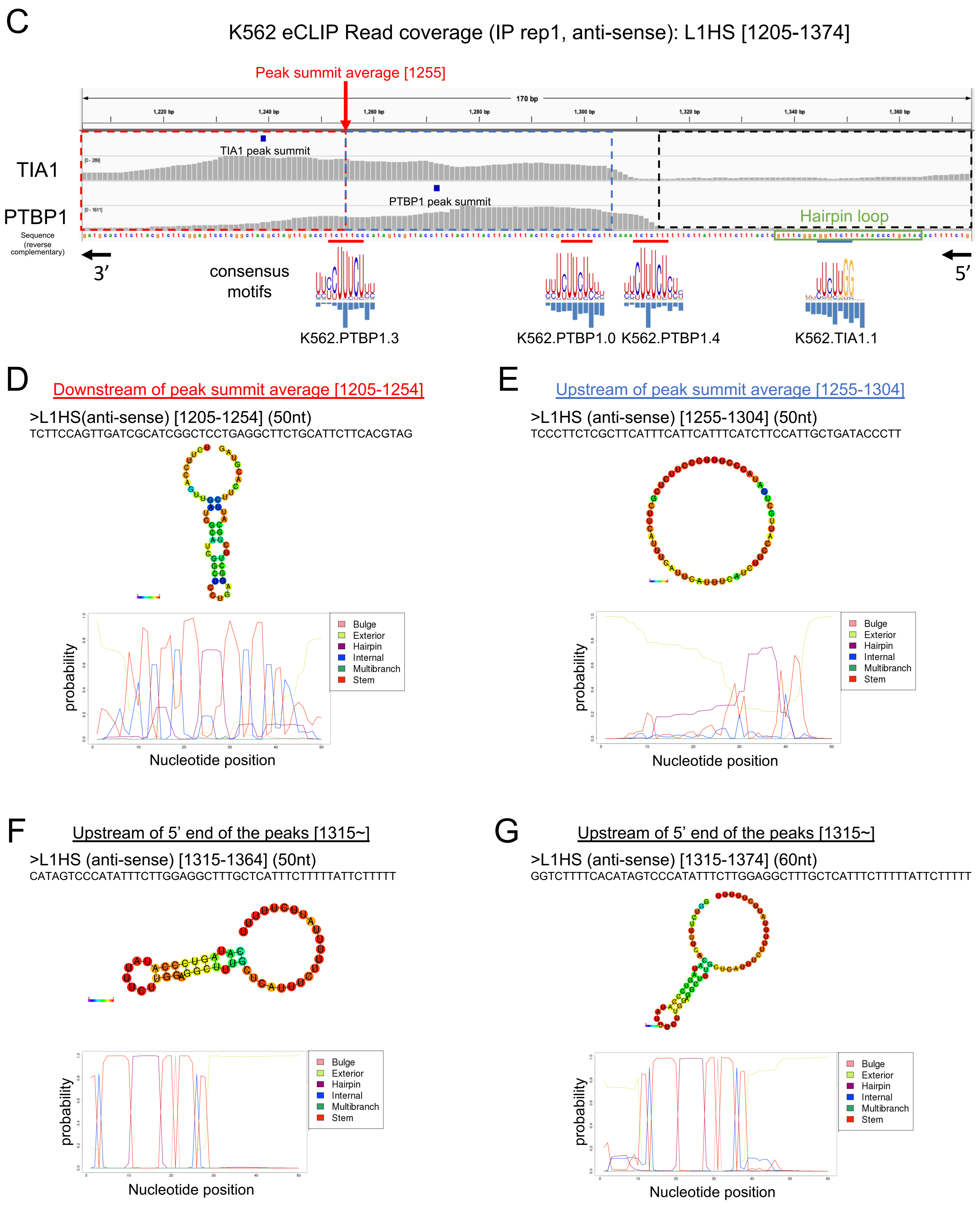

Supplementary Figure 5

(C) eCLIP IP read coverage of RBPs associated with L1HS[1205-1374](anti-sense) in K562 cells. The y-axis shows the names of RBPs, and the x-axis shows the nucleotide position of the L1HS sequence. The red or blue rectangles indicate the upstream or downstream regions of the peak summit average [1255] that were found to form the RNA secondary structures shown in (D), respectively. The black rectangles indicate the upstream region of the 5' end of the peaks [1315~] that was found to form RNA secondary structures shown in (E).

(D, E, F, G) Prediction of RNA secondary structures of anti-sense L1HS for the downstream region of the peak summit average [1205-1254](D) or the upstream region of the peak summit average [1255-1304](E) or 50 nt (F) or 60 nt (G) upstream regions of the 5' end of the peaks [1315] calculated using CentroidFold (upper panels) or CapR (lower panels). Colored bases of the upper panels (CentroidFold) indicate the base-pairing and loop probabilities. The lower panels (CapR) show the structural profile of an RNA base for a set of six probabilities (stem part, hairpin loop, bulge loop, internal loop, multibranch loop, and exterior loop) that the base belongs to each category. The x-axis indicates the nucleotide position, and the y-axis indicates the probability of the structural profiles.

Supplementary Figure 6

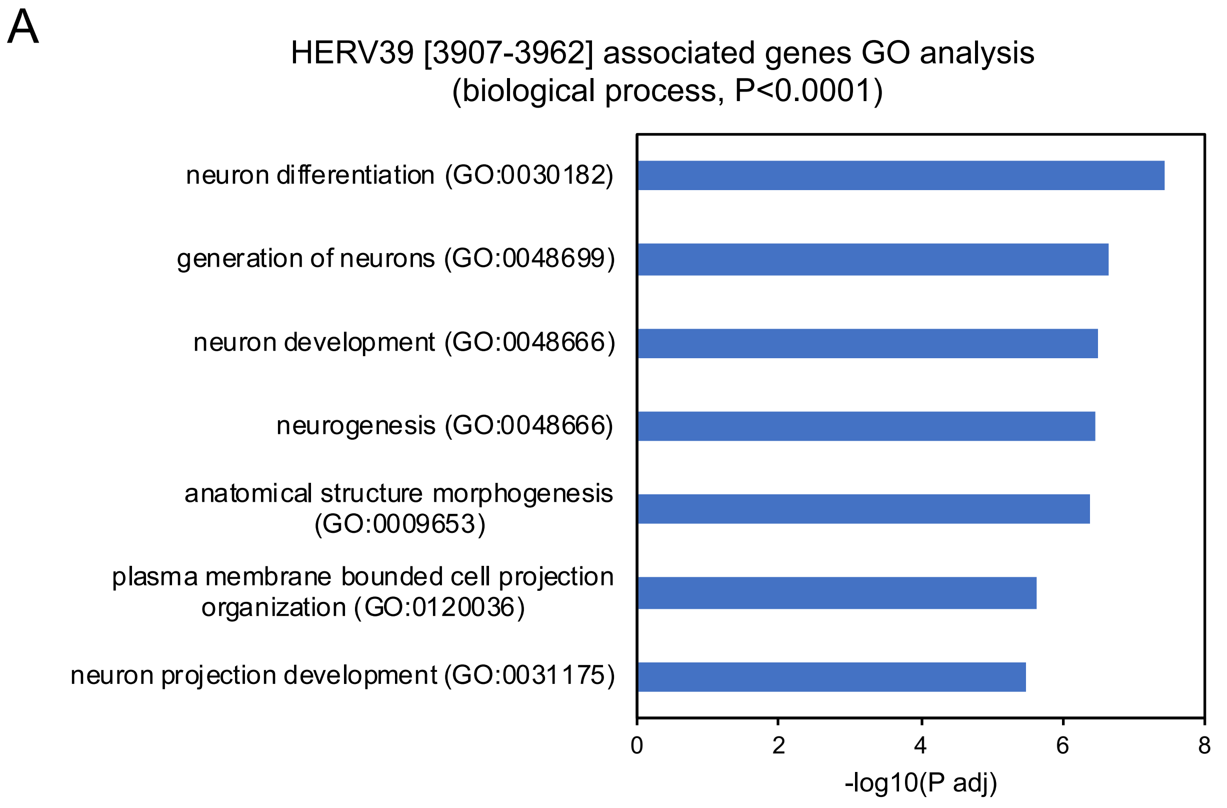


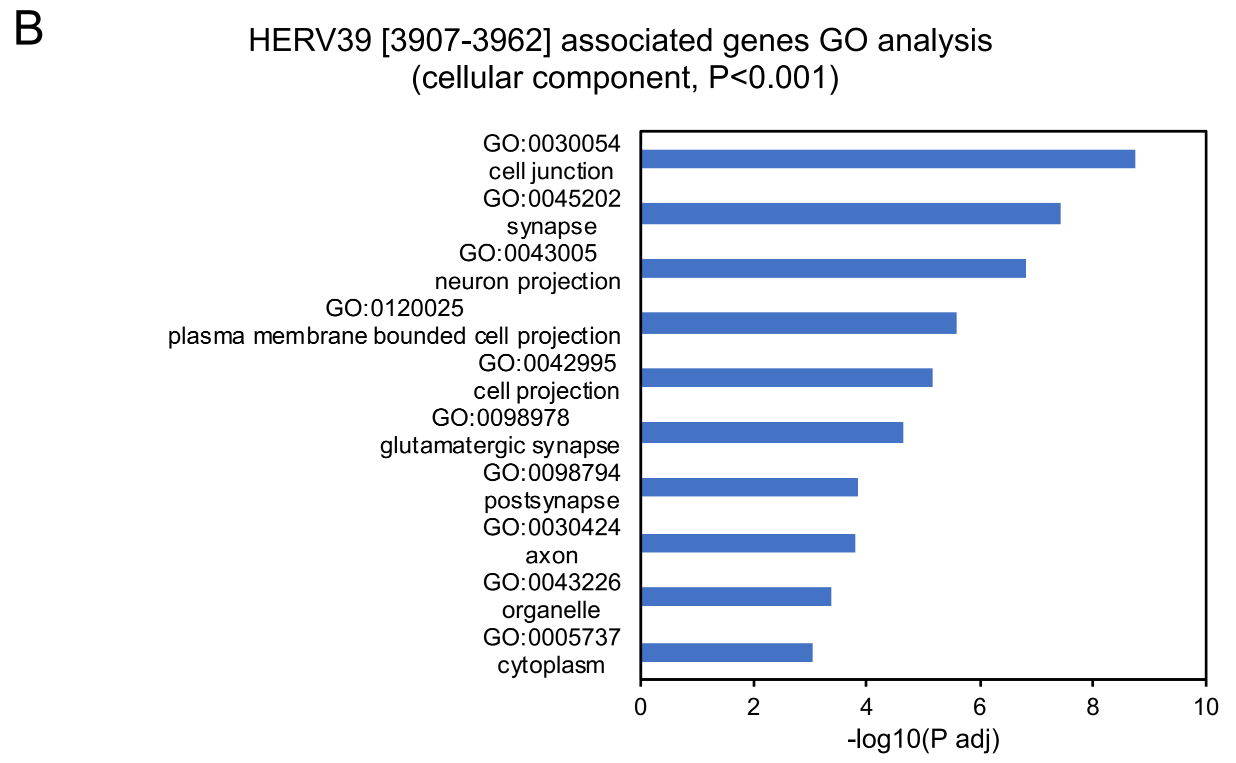


Supplementary Figure 6
(A,B) Gene ontology analysis of HERV39 [3907-3962] associated genes for biological process (A) or cellular component (B). The x-axis indicates the -log10 (adjusted p-value).

Supplementary Figure 7

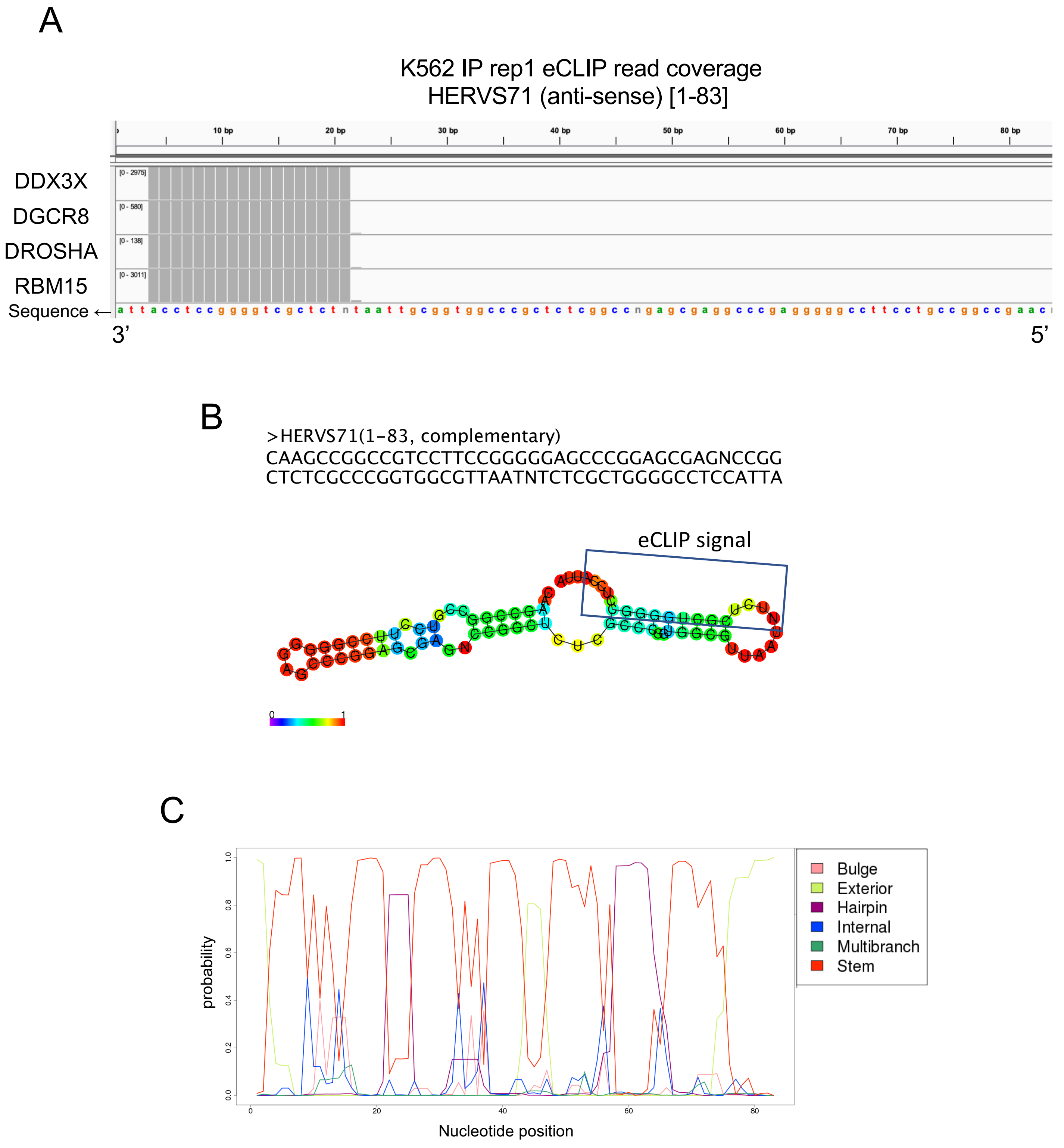


Supplementary Figure 7
(A) eCLIP IP read coverage of RBPs associated with HERVS71[1-83] (anti-sense) in K562 cells. The y-axis shows the names of RBPs, and the x-axis indicates the nucleotide position of the HERVS71 sequence. (B, C) Prediction of the RNA secondary structures of anti-sense HERVS71 for the upstream and downstream region of the 5' end of the peaks [21] calculated using CentroidFold (B) or CapR (C). (B) The blue rectangle indicates an eCLIP peak signal. Colored bases indicate the base-pairing and loop probabilities. (C) The structural profile of an RNA base for a set of six probabilities (stem part, hairpin loop, bulge loop, internal loop, multibranch loop, and exterior loop) that the base belongs to each category. The x-axis indicates the nucleotide position, and the y-axis indicates the probability of the structural profiles.
