## Supplemental Materials and Methods for "Binding patterns of RNA binding proteins to repeat-derived RNA sequences reveal putative functional RNA elements"

Supplementary Materials and Methods

Cutadapt:

Round1:

cutadapt -f fastq --match-read-wildcards --times 1 -e 0.15 -O 7 (11 if adapter=A03, G07, A04, F05) --quality-cutoff 6 -m 18 -g(adapter)

Round2:

cutadapt -f fastq --match-read-wildcards --times 1 -e 0.1 -O 1 --quality-cutoff 6 -m 18 -a(adapter) -A(adapter)

Round3:

cutadapt -f fastq --match-read-wildcards --times 1 -e 0.1 -O 5 (9 if adapter=A03, G07, A04, F05) --quality-cutoff 6 -m 18 -A(adapter)

adapter sequences:

| adapter_name | Read1_5’ | Read2_3’ |
|---|---|---|
| A01 | CTTCCGATCTAAGCAAT | ATTGCTTAGATCGGAAGAGCGTCGTGT |
| B06 | CTTCCGATCTGGCTTGT | ACAAGCCAGATCGGAAGAGCGTCGTGT |
| C01 | CTTCCGATCTACAAGTT | AACTTGTAGATCGGAAGAGCGTCGTGT |
| D08 | CTTCCGATCTTGGTCCT | AGGACCAAGATCGGAAGAGCGTCGTGT |
| A03 | CTTCCGATCTATGACCNNNNT | ANNNNGGTCATAGATCGGAAGAGCGTCGTGT |
| G07 | CTTCCGATCTTCCTGTNNNNT | ANNNNACAGGAAGATCGGAAGAGCGTCGTGT |
| A04 | CTTCCGATCTCAGCTTNNNNT | ANNNNAAGCTGAGATCGGAAGAGCGTCGTGT |
| F05 | CTTCCGATCTGGATACNNNNT | ANNNNGTATCCAGATCGGAAGAGCGTCGTGT |
| X1A | CTTCCGATCTNNNNNCCTATAT | ATATAGGNNNNNAGATCGGAAGAGCGTCGTGTAG |
| X1B | CTTCCGATCTNNNNNTGCTATT | AATAGCANNNNNAGATCGGAAGAGCGTCGTGTAG |
| X2A | CTTCCGATCTNNNNNTATACTT | AAGTATANNNNNAGATCGGAAGAGCGTCGTGTAG |
| X2B | CTTCCGATCTNNNNNATCTTCT | AGAAGATNNNNNAGATCGGAAGAGCGTCGTGTAG |
| RiL | CGACGCTCTTCCGATCT | AGATCGGAAGAGCGTCGTGT |

STAR

STAR --runMode alignReads --runThreadN 32 --genomeDir /home/ /ENCODE/Repbase-Human --readFilesIn /home/ENCODE/eclip-data/ /K562_???_IP_repX_R1.adapterTrim.round3.rmDup.fastq.gz /home/ENCODE/eclip-data/K562_???_IP_repX_R2.adapterTrim.round3.rmDup.fastq.gz --outSAMunmapped Within --outFilterMultimapNmax 1 --outFilterMultimapScoreRange 1 --outFileNamePrefix /home/ENCODE/K562_???_IP_repX --outSAMattributes All --readFilesCommand zcat --outStd BAM_Unsorted --outSAMtype BAM Unsorted --outFilterType BySJout --outReadsUnmapped Fastx --outFilterScoreMin 10 --outSAMattrRGline ID:foo --alignEndsType EndToEnd > /home/ENCODE/ /K562_???_IP_repX.uniq.bam

Peak call

Piranha -s -b 60
